# Coexistence of state, choice, and sensory integration coding in barrel cortex LII/III

**DOI:** 10.1101/2023.04.10.536289

**Authors:** Pierre-Marie Gardères, Sébastien Le Gal, Charly Rousseau, Alexandre Mamane, Dan Alin Ganea, Florent Haiss

## Abstract

During perceptually guided decisions, correlates of choice are found as upstream as in the primary sensory areas. However, how well these choice signals align with early sensory representations, a prerequisite for their interpretation as feedforward substrates of perception, remains an open question. We designed a two alternative forced choice task (2AFC) in which mice compared stimulation frequencies applied to two adjacent vibrissae. The optogenetic silencing of individual columns in the primary somatosensory cortex (wS1) resulted in predicted shifts of psychometric functions, demonstrating that perception depends on focal, early sensory representations. Functional imaging of layer II/III single neurons revealed sensory, choice and engagement coding. From trial to trial, these three varied substantially, but independently from one another. Thus, coding of sensory and non-sensory variables co-exist in orthogonal subspace of the population activity, suggesting that perceptual variability does not originate from wS1 but rather from state or choice fluctuations in downstream areas.

## Introduction

The brain guides the body by processing incoming sensory information and using it to select contextually relevant actions. This perceptually dependent process has been shown to involve at least two components: the encoding of sensory evidence, transiently elicited in sensory areas, and the progressive accumulation of motor variables in a distributed network of decisional and motor areas (Gold and Shadlen 2007; Hernández et al. 2010; Siegel et al. 2015). To understand where and how sensory representations inform decisional processes, it is necessary to identify features of the neuronal activity that encode simultaneously sensory and choice information.

Pioneering studies addressed that issue using two alternative forced choice tasks (2-AFC), where primates had to judge the motion direction of a cloud of points (Britten et al. 1992; Salzman et al. 1990; Britten et al. 1996). Perception was reported by choosing one of two possible actions. These studies found that trial-by-trial fluctuations of neuronal activity in the visual areas predicted choices of the animal, i.e., display choice related activity (Britten et al. 1996). The firing rate of neurons in sensory areas is hence thought to carry the code for sensory representation informing behavior. In the somatosensory system of the primate, similar approaches have revealed that choice related activity arises from the secondary somatosensory area and higher in the decisional hierarchy (Romo et al. 2002; Hernández et al. 2010) . More recently, research in rodents has identified choice related activity forming already at the level of the primary somatosensory area (S1) (Sachidhanandam et al. 2013; Yang et al. 2016), It is yet difficult to conclude that the choice signals in S1 have a causal influence on behavior and perception.

First, many studies in the rodent model used a go/nogo paradigm and studied the detection of a stimulus close to the detection threshold. The widespread cortical activity related to onset of facial movement (Musall et al. 2019; Stringer et al. 2019)– including in sensory areas– renders ambiguous the signals associated with go trials. These approaches are also relatively vulnerable to animal biases and changes in motivation. It has thus been proposed that more complex designs, such as 2AFC with delayed response are needed to disentangle choice from these other sources of modulation (Zagha et al. 2022).

Perhaps more important, choice signals are generally described as increases in activity indiscriminately in a fraction of sensory or non-sensory neurons. If a sensory representation is used to inform the behavior, its fluctuation from trial to trial should have an impact on the choice. In other terms, the sensory code, rather than the activity in the area, should co-fluctuate with the behavior (Panzeri et al. 2017). Sensory and choice coding in the rodent S1 has been studied mostly separately, or at the level of single neurons. Therefore, a description of how choice versus early sensory representation are differentially organized in S1 neuronal populations is lacking.

To tackle these challenges, the focal and quasi-discrete sensory representation of single whiskers in S1 represents a considerable advantage. Here we developed a novel two alternative forced choice task that would allow us to monitor and manipulate the representations of the two competing sensory alternatives.

## Results

### Discrimination behavior of vibro-tactile frequencies applied to adjacent whiskers

Mice were water deprived and required to compare two frequencies (F1 and F2) delivered simultaneously to two adjacent whiskers on the same row of the whisker pad (e.g. B1/C1). The two target whiskers, designated thereafter as W1 and W2, were respectively associated with left and right waterspouts, designated thereafter as choice 1 and choice 2 (Fig. 1a). Tactile stimulation lasted 1 second after which waterspouts came in a reachable position. At that time, the mice were allowed to lick during a two second decision period (Fig. 1b). A water reward was delivered if the subject first licked the spout associated with the whisker deflected at the highest frequency. Importantly in this task, the frequencies are proportional to the average speed of whisker deflection and thus directly represent intensity of the stimulation (Waiblinger et al. 2015). While the subject was performing the task, we recorded videos of its face and snout, allowing us to track multiple behavioral variables such as the pupil size, movement from the whisker, nose, and tongue (Fig. 1b; see Methods).

**Figure 1.**
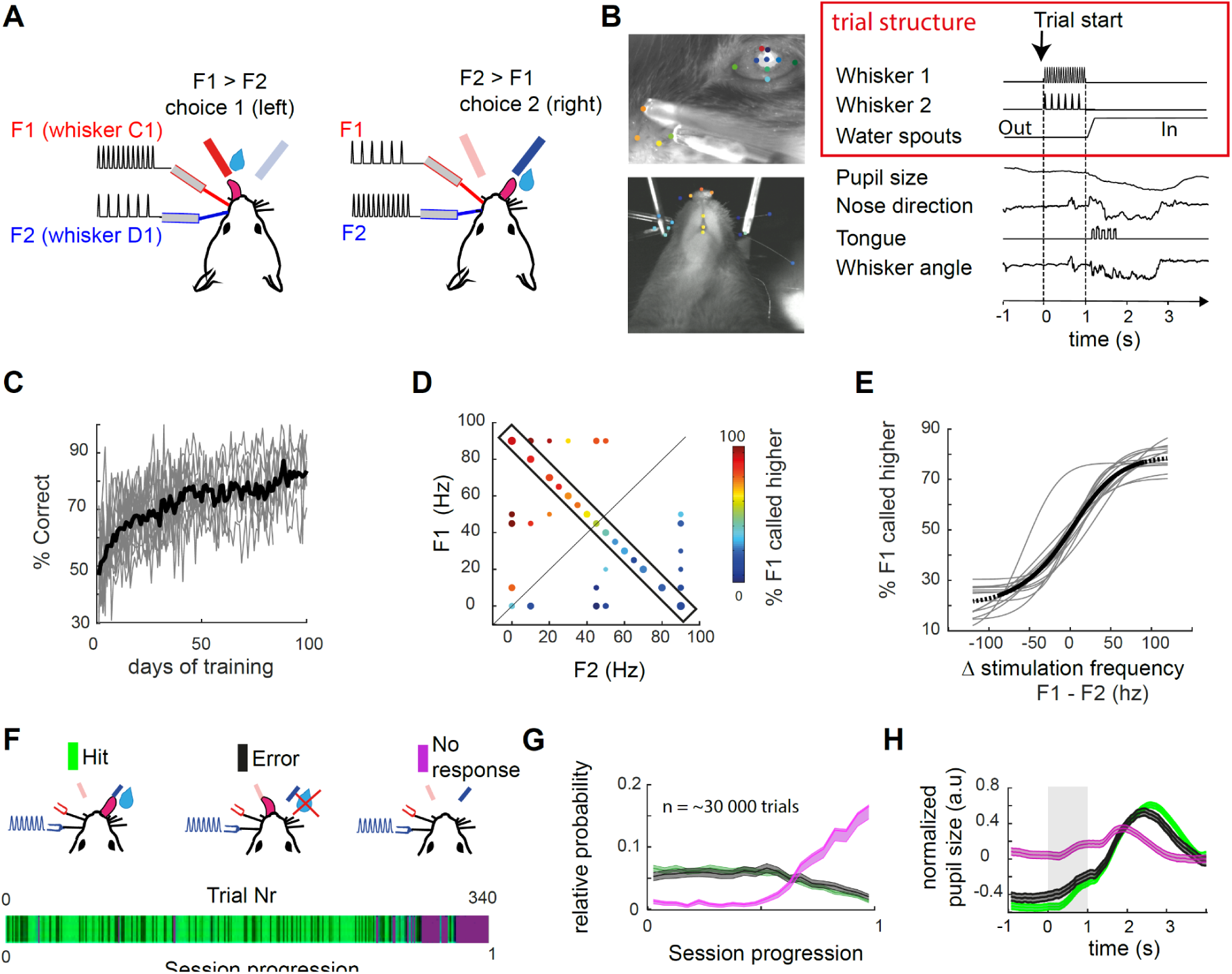
Performance and behavior during discrimination of simultaneous vibrotactile stimulation applied to adjacent whiskers. A, Schematic of the task. left: W1 whisker is deflected with higher frequency than W2, left licks trigger a reward delivery. Right: F2>F1, licks to the right trigger a reward delivery. B, Face tracking and trial structure. Left: two example views simultaneously recorded. Some face parts elements are tracked using DeepLabCut (Mathis et al. 2018) and here labelled with random colors (see methods). Right: trial time structure with example face part tracking (downsampled to 30 Hz). Waterspouts come in a reachable position at the end of the whisker stimulation. Scales are normalized between minimum and maximum amplitudes C, Learning curves of 15 individual mice. Calcium imaging and further analysis were carried on a subset of sessions where the animal is considered expert (>70% Correct trials). D, Behavioral performance in the stimulus space. Each point represents a set of two frequencies applied to the two whiskers. The diagonal represents the task boundary (F1 = F2). Proportion of left/right choices are color coded. E, Psychometric function drawn from conditions within the rectangle in D. (i.e. with a constant summed frequency F1 + F2 = 90 Hz) circles represent average performance and thin traces are logistic fits for individual animals (see methods). Black thick line represents a logistic fit for the pooled dataset. F, Three possible behavioral outcomes in one example session. Top: 3 possible behavioral outcomes: first lick right, first lick left, and no lick during the response window. Bottom: example session with behavioral outcome color coded. The trial index is a normalization of trial number between 0 and 1 (respectively first to last trial of the session). G, Relative distribution of hits, errors, and misses within the daily session progression (average across sessions and mice), normalized by category. Misses typically appeared in blocks at the end of the session. H, Normalized pupil size in the pre-stimulus period predicts engagement of the subject. Error bars / error shades represent s.e.m.

The initial training phase for the 2AFC started with single whisker stimulation only and subjects were trained to respect the delay to the end of the stimulus (Fig. 1c). Once they reliably scored 70% in that task, F1 and F2 were set to vary in a range of frequencies going from 0 to 90 Hz, defining the stimulus space (Fig. 1d). To quantify the relationship between the stimulus and the sensory capabilities of mice, we fitted the probability of F1 being called higher with a psychometric function (Wichmann and Hill 2001) which reveals a reliable dependence of the choice side on the stimulation frequency difference ΔF = F1 - F2 (Fig. 1e) across animals. ΔF is the dimension explaining most performance across the stimulus space (Fig. S1). Importantly, mice were able to compare the frequencies at all points tested in the stimulus space (Fig. 1d). The same stimulus (e.g. F1 =50 Hz) could be contextually categorized as a distractor or target, excluding that the task is solved based on the recognition of a specific target stimulus only (Hernández et al. 1997). Instead, we conclude that mice were capable of generalizing the abstract rule of frequency comparison across the entire stimulus space.

To distinguish the different behavioral outcomes, we used the animal’s licking behavior and its facial movements. The 2AFC task design leads to three possible trial outcomes: correct trials, error trials, and no response, when no spouts are licked during the decision period (Fig. 1g). Most of the no-response trials were recorded in blocks at the end of the daily session, so we hypothesized a different state of task disengagement. This view is strengthened by a physiological marker, as we observe larger pupillary dilation in the baseline epoch and weaker pupillary response to stimulation, consistent with previous report (baseline dilation p= 0.036, difference between Correct and Miss, n=4 mice, Friedmann test with tuckey’s post hoc correction; Fig. 1h)(Ganea et al. 2020). We found that within the engaged state, nose movements are the most indicative of the animal’s earliest reaction and decision (Fig. S2). We thus used their onset as a proxy for the first movement executed by the animal in each trial. Accordingly, we distinguished three trial categories: first, trials with movements prior to stimulus onset (24.4 ± 3.1 % trials; mean ± s.e.m across animals, n = 7 mice.) which are excluded from further analysis; second, “impulsive” trials with movements prior to the end of the delay (41.2 ± 2.9 % trials), and third, trials in which the animals refrain from moving until the decision period (34.4 ± 5.4 % of the trials, no significant movement of the snout, tongue or whiskers, Fig. S2). Therefore, the delayed 2AFC task design with video tracking enables distinction of engaged versus disengaged states, and enables separation of motor from choice variables in a subset of trials with controlled delayed responses.

### Somatotopic whisker representation and increased activity level with stimulation frequencies

To characterize S1 sensory representations, we used two photon calcium imaging guided by intrinsic optical imaging (IOI, Fig. 2a-c, see Methods), recording hundreds of L2/3 neurons simultaneously in the same field of view (337 ± 20 neurons per FOV). Transient fluorescence responses evoked by single whisker deflections matched the spatial arrangement of IOI maximum intensity (Fig. 2c). We considered the activity of single neurons in response to single and dual whisker stimulation frequencies. Fluorescence traces were detrended, and firing rate (FR) was inferred using a deconvolution algorithm (see Methods; Fig. 2b). Consistent with previous studies (O’Connor et al. 2010; Mayrhofer et al. 2015; Barz et al. 2021), we found that a small fraction of neurons were highly responsive to tactile stimulation (∼10 %; Fig. 2c). Yet, a larger fraction of the population had significant response compared to baseline: 32.3 ± 4.8 % of neurons were suppressed and 48.1 ± 4.7 % of neurons were activated, mean ± s.e.m across n= 11 FOV, all trials included. Neurons responding to one whisker showed increased probability to also respond to the other whisker (r² = 0.49; similarly to previous reports)(Clancy et al. 2015), but paradoxically the fraction of most responsive neurons (∼10%;) displayed the highest whisker selectivity index (SI) and were matching more closely the somatotopic map (Fig. 2d, Fig. S3). In a separate set of animals with conditional genetic expression of GFP in GABAergic neurons (VGAT-Cre), we quantified selectivity and spatial distribution of labeled inhibitory neurons (INs) and putative excitatory neurons (ENs)(see methods; Fig. 2e). IN population selectivity indexes were lower than the EN population, but ranged on a similar scale (p<0.001, |µSI_IC_|=∼ 0.33, |µSI_EC_ |=∼ 0.40 Mann-Whitney U test, Fig. 2e) INs selectivity was also somatotopically distributed (Fig. 2e). These results highlight the functional selectivity of inhibitory and excitatory neurons in two neighboring microcircuits. The spatial transition of functional selectivity illustrates the somatotopic organization into cortical columns.

**Figure 2.**
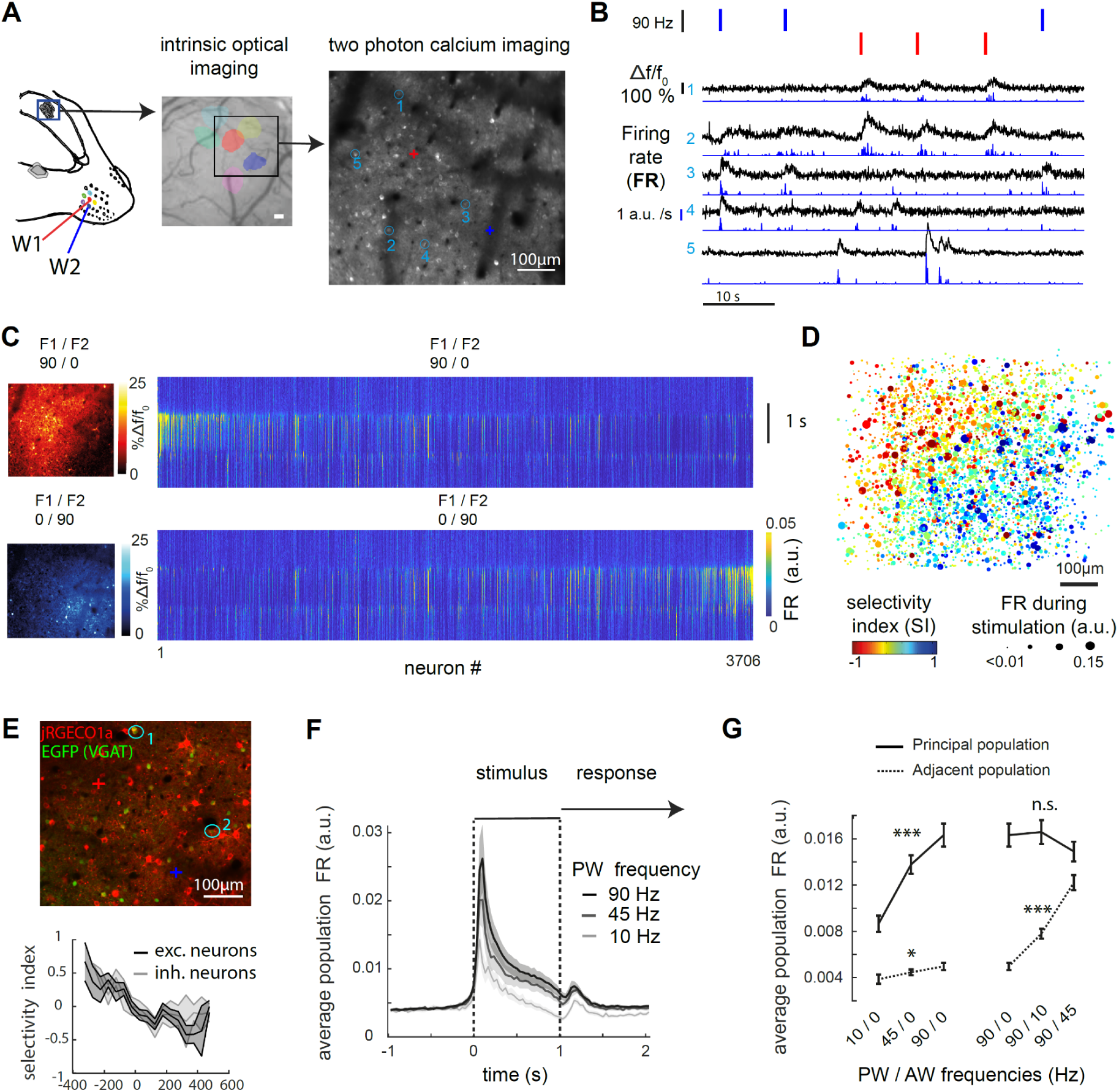
Activity levels in neighboring cortical columns increase with stimulation frequencies of their preferred whisker. A, Simultaneous Imaging of W1 and W2 cortical representations in wS1 L2/3. Left: Contralateral recording in the head-fixed mouse. Middle: Example Intrinsic optical imaging. Right: Example Ca2+ imaging field of view (FOV) spanning the two barrels; blue circles indicate example ROIs for which calcium transients are depicted in (D). B, Example of fluorescence and deconvolution traces for five example neurons over six consecutive trials. Top colored lines represent tactile stimulation of W1 only (red) or W2 only (blue). Black Traces: Neuronal calcium transients, depicted as Δf/f0. Blue traces: Firing rate (FR) in arbitrary units (a.u.) resulting from the deconvolution. C, Response to single whisker stimulation at 90 Hz (W1 and W2, top and bottom rows respectively). Left: pixel-wise change in fluorescence Δf/f0 is color coded. Right, average FR of each neuron in response to single whisker stimuli. Neurons are sorted by increasing FR_1_ - FR_2_, with FR_1_ and FR_2_ representing firing rate during 90 Hz stimulation of W1 and W2. Same sorting order in top and bottom plots. D, All neurons from 11 FOVs aligned to the mid-points between barrels. Selectivity index (SI) is represented by color and average FR is represented as dot size. FR= FR_1_ + FR_2_; SI = (FR_1_ + FR_2_)/ (FR_1_ - FR_2_). E, Inhibitory versus excitatory response to whisker stimulation. Top: Example FOV of a VGAT-cre mice, Barrel centers are marked by a cross. Bottom: average SI for INs and putative ENs (n = 267 and 974 neurons respectively from 6 FOV in 2 animals). SI averaged in bins of 50 µm. Error shades represent CI95. F, Population average FR in response to three stimulation frequencies of single whisker stimulation. Error shades represent s.e.m. across FOVs, n = 11. G, Average FR of the principal whisker (PW) population and adjacent whisker (AW) population in response to single and multi-whisker stimulation. Statistical test represents increase or decrease of population activity with the stimulation frequency of the PW (left) or AW while PW is held to 90 Hz stimulation (right). *** p<0.001 with Spearman correlation test. Error bars represent s.e.m. across FOVs.

To understand how stimulation frequencies influence these microcircuits’ response level, we split the neuronal population in two, based on their preferred whisker (SI > 0 and SI < 0). In response to their preferred whiskers stimulation at 90 Hz, the a sub-populations instantaneous firing rate increases by a factor 3.3 immediately after stimulus onset, and then adapted but remained over baseline activity level (increased by factor 1.8 ± 0.2, mean ± s.e.m., 0.5 to 1 s after stimulus onset, n = 11 FOVs; Fig. 2f). Importantly, in response to increasing frequencies, we observe an increase in the average population activity (Fig. 2f-g, principal population). Adjacent populations respond positively but weakly to the stimulation of the principal whisker alone. Finally, adding increasing adjacent whisker’s stimulation over principal whisker’s stimulation held constant, seemed to lower the response of the principal whisker’s population (Fig. 2g), suggesting a suppressive effect of multi-whisker stimulation. As frequencies are directly proportional to mean speed deflection of the whisker, frequencies also represent an intensity of stimulation (Waiblinger et al. 2015). In accordance with this concept, our data show that increasing frequencies are represented by monotonical increase in activity level in the principal whisker column.

### Neural activity in a single wS1 cortical column drives perception of the whisker stimulation intensity

As the two sensory alternatives have distinct spatial representations, we evaluated the possibility of manipulating them selectively. Optogenetic wild-field stimulation was combined with two-photon functional imaging (Fig. 3a; see methods)(Prsa et al. 2017). The efficiency and selectivity of optogenetic stimulation was surveyed in four distinct conditions: no light, broad illumination of the two barrels and selective stimulation of either single barrel (Fig. 3b-c). Across layers of the neocortex, light was shaped in a cylindrical manner (Fig. S4). Maximum light intensity stimulation (17.9 mW.mm² with 450 µm light disk, as measured below the objective) during epochs of spontaneous activity induced a significant increase in activity for 23.2% cells, being putative IN, and a significant decrease for 20.8% cells, being putative EN (p <0.01, t-test versus spontaneous activity across trials). These proportions were dependent on the distance to the illumination center (Fig. S4). We then explored how optogenetic stimulation of a single barrel INs influences the sensory signals associated with single whisker deflection (Fig. 3d). At all light powers, optogenetic stimulation decreased sensory response to the preferred whisker in both target and neighboring columns. But the reduction of sensory activity was always stronger for the optogenetically targeted barrel. Taken together these results indicate that it is possible to differentially manipulate the two sub-ensembles, shifting bi-directionally the ratio of response toward one or the other whisker representation.

**Figure 3.**
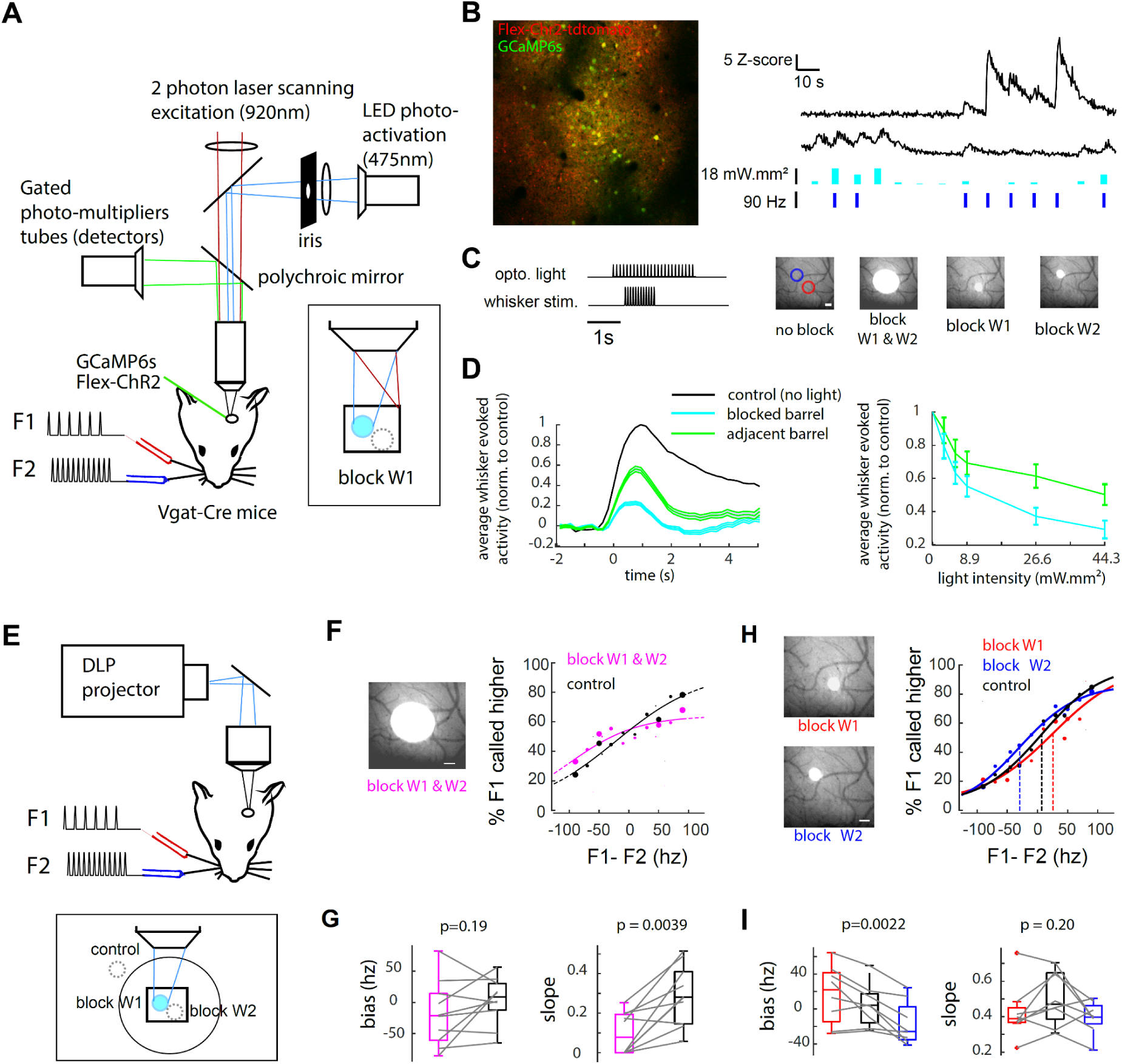
Differential optogenetic inhibition of single barrel induces selective shifts in perception. A, Setup for combined optogenetic and calcium imaging. A fast gated PMT and a polychromatic mirror allowed the use of blue light interleaved with two-photon calcium imaging. B, Left: Example FOV of one VGAT-cre animal following injection of GCaMP6s and Cre-dependant ChR2. Right: example fluorescence transients in response to whisker and light stimulation for a putative excitatory neuron (top) and a putative inhibitory neuron (bottom). Light and whisker stimulation amplitude are indicated below. C, Optogenetic stimulation parameters. Left: Trial structure. 1ms pulses of light were delivered at 40 Hz during 2.5 s. Right: Different structured light patterns, from left to right: - W1 and W2 barrels are illustrated as colored circles centered on IOI max response (200 µm diameter); - Illumination of the two barrels (450 µm diameter disk, FWHM); - selective illumination of barrel W1 (105 µm diameter disk, FWHM); - selective illumination of barrel W2. D, Impact of selective inhibition (105 µm diameter disk) onto targeted and adjacent barrels evoked response (n = 6 barrels in 3 animals with 503 neurons in total). Inhibition is measured as the residual evoked response elicited by the preferred whisker 90 Hz stimulation with/without optogenetic light. Inhibitory neurons and non-whisker responsive neurons were excluded from the analysis (see details in the methods). Error bars/shades represent 95% confidence intervals. Left: time course of whisker evoked response. Right: mean residual activity across different optogenetic light intensities. E, Setup for optogenetic during behavior. A DLP projector was used to target barrels with 1 ms pulse at 40 Hz with light power ranging between 8.9 and 26.6 mW/mm² F, Non-selective barrel inhibition. Left: example FOV inhibition of the barrels, right: Psychometric fit during intermingled trials with either sham optogenetic (black) and 450 µm diameter disk optogenetic blocking (magenta). G, Psychometric fit difference, compared in the same mice across optogenetic conditions. Same color code as for (F). H, Selective barrel inhibition. Left: example FOV inhibition of the barrels, right: psychometric fit during intermingled trials with either inhibition of W1 barrel (red), inhibition of W2 barrel (blue) or sham control (black). I, Psychometric fit difference, compared in the same mice across optogenetic conditions. Same color code as for (H). In F-I, we included the same number of trials in each optogenetic and control condition. Trials were paired with closest occurring sham control trials (i.e. therefore we ensured that only intermingled trials were included, to avoid bias due to repetitive optogenetic presentation). Trials were matched for all statistical analysis and psychometric fits.

To identify the involvement of S1 in the sensory guided decision, trials with and without optogenetic stimulation were interleaved during the behavioral task (Fig. 3 e-i). Optogenetic stimulation was provided in 25% of the trials, versus a control condition in which light was pointed on the dental cement next to the brain (Fig. 3e). In response to the broad stimulation pattern, we found that the behavioral discrimination was significantly impaired in all stimulation conditions as compared to the control condition (slope of the psychometric fit; p = 0.0039, n= 10 mice; Fig. 3f-g; Wilcoxon sign-rank test). This result suggests that wS1 is causally involved in the perceptual decision making-task used here. However, the perturbation of activity homeostasis at a larger scale -including other brain areas-(Otchy et al. 2015), might potentially be responsible for the impairment of behavior, independent of the manipulation of the sensory representation.

To test whether single whisker column manipulation selectively promotes one choice versus the other, we used the selective inhibition of barrels corresponding to whisker W1 or whisker W2 (Fig. 3h-i) intermingled with the control condition. The single column inhibition induced a bidirectional behavioral bias in the psychometric curve depending on the silenced barrel (psychometric bias, p =0.0022, Friedman test, n= 8 mice; Fig. 3i). This horizontal shift in the psychometric curve gives us indications on the relationship between S1 activity and subjective report. At the used light intensities, optogenetic inhibition of column W1 induces on average a rightward shift of the curve of 13.4 Hz ± 6.5 (mean ± s.e.m.), and inhibition of column W2 a leftward shift of 26.4 Hz ± 5.9 (mean ± s.e.m.). To control whether the optogenetic intervention alters the motor response instead of the sensory perception, we compared proportion of choice 1 in trials with optogenetic but without whisker stimulation and found it similar, although slightly biased to ipsilateral choices (fraction of choice 1, W1 barrel blocking: 0.559 ±0.043; W2 barrel blocking: 0.522 ± 0.034, mean ± s.e.m. p = 0.21, Wilcoxon signed rank test, n=7 mice). Thus, despite a seemingly modest amplitude, the bidirectional bias provides a strong level of evidence for the causal influence of single column activity on the perceived whisker stimulation intensity.

### Multi-whisker suppression enables generalization of the frequency comparison task in the stimulus space

To further dissect single neuron tuning to the whisker stimulation parameters, we applied a multiple linear regression (Fig. 4a)(Romo et al., 2002):

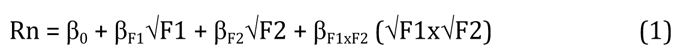

Where three parameters of whisker stimulation best explain the activity of neurons. First and second are the square roots of single whisker stimulation frequencies (F1 and F2; Fig. 4a). Third, the supra-linear interaction of the two whiskers frequencies F1xF2 increases the model fit (i.e. lower Akaike criterion and higher explained variance; Fig. S5). Other regressions parameters were tested, such as divisive interaction terms or full categorical regressor (i.e. modal tuning to single frequencies). These could only marginally improve the overall model fit of single neuron activity at the cost of more complex designs, and were thus excluded. Overall, only a fraction of single neurons’ activity variance could be explained by the stimulation parameters (r2= 8.1 ± 0.2% mean ± s.e.m.; n = 3706 neurons; Fig. 4a), in agreement with previous reports (Musall et al. 2019). We observe across the S1 population a positive correlation between β_F1_ and β_F2_ weights (Fig. 4b, r = 0.55; Pearson correlation between β1 and β2) meaning that neurons respond to the two whiskers and that representations of the whiskers are overlapping in the population. However, the more the neurons are driven by single whisker frequencies, the more they are being suppressed by the interaction of the two whiskers (Fig. 4b, r = -0.65, r = -0.62; Pearson’s correlation between β_F1_ and β_F1xF2_, and between β_F2_ and β_F1xF2_, respectively, p<0.001). This regression analysis confirms that single neuron’s response increases with stimulation frequency, and shows that multi-whisker suppression is an important factor of L2/3 neuronal activity.

Does multi-whisker suppression help solving the comparison task? To evaluate how this computation affects activity level in the stimulus space, three example neurons were compared: One with multi-whisker suppression, one without multi-whisker integration and one with multi-whisker supralinear responses (neurons 1, 2 and 3 respectively in Fig. 4c-e). The neuron with multiple whisker enhancement shows no difference in activity level on either side of the boundary task. Therefore, its activity level is not informative in regard to the task, and may rather represent a source of noise in our paradigm. On the contrary, multi-whisker suppression leads the gradient of activity in the stimulus space to become perpendicular to the task boundary. Activity level reflects the difference in stimulation frequency F1-F2 rather than the stimulation frequency of their preferred whisker (Fig. 4e). Therefore, the output of this neuron encodes the stimuli in a relevant way for the generalization of frequency discrimination, as compared to the neuron without suppression. This comparison reveals how multi-whisker suppression increases the discriminability of neighboring stimulation frequencies

**Figure 4.**
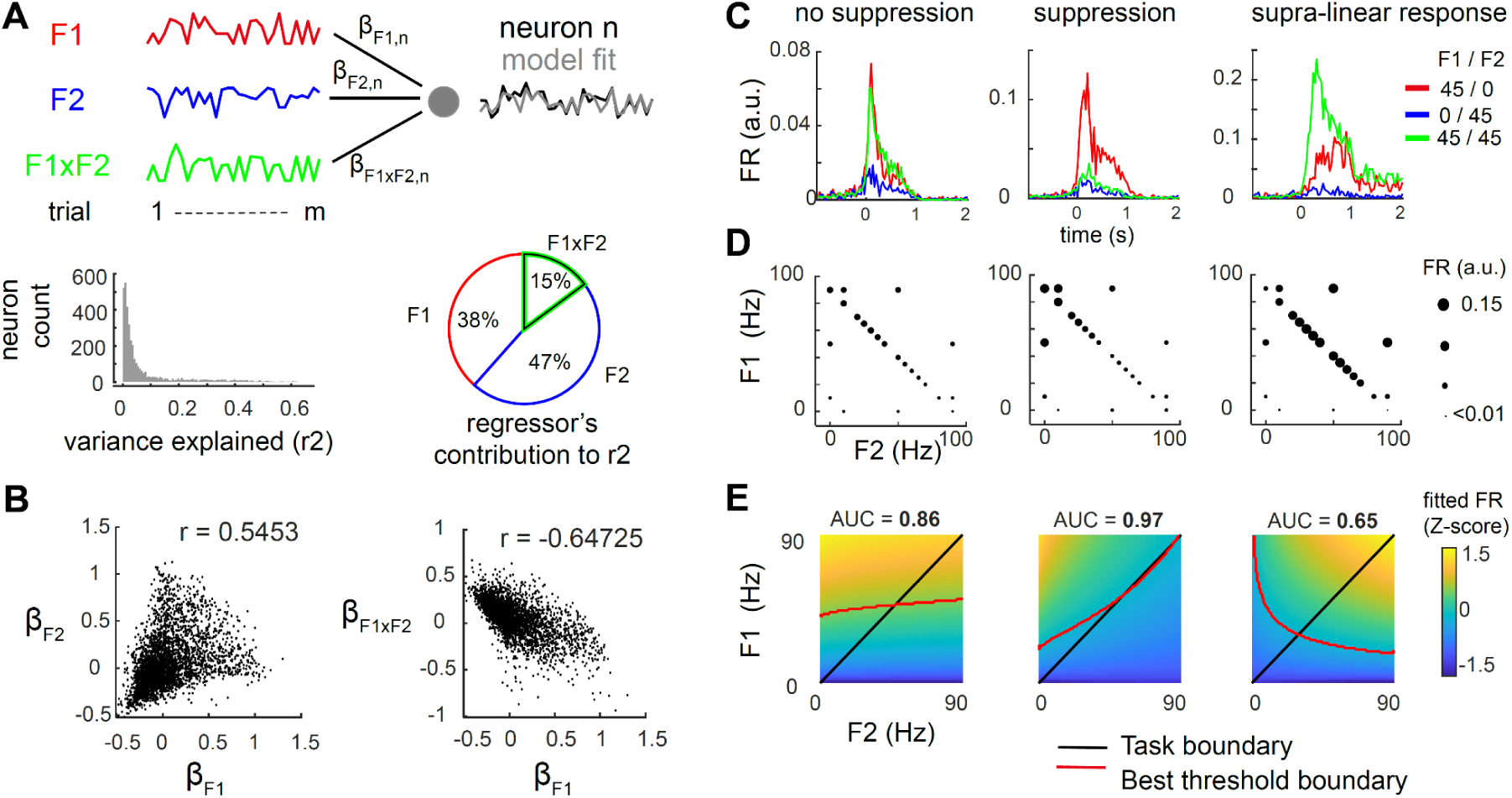
Multi-whisker suppression improves frequency comparison performance in the stimulus space. A, Linear regression of single neuron activity on whisker stimulation frequencies. Top row : Example regression of one neuron in 20 trials (see text and methods). Bottom left: fraction of explained variance across neurons. Bottom right: fraction of variance explained by each regressor (average over n= 3706 neurons), relative to the total variance explained by the model. B, Cell by cell distribution of model weight. Note the positive correlation between single whisker weights (left) and negative correlation between single and interaction weights (right). C-E, Three example neurons (left to right columns) showing different forms of multi-whisker integration. C, Average firing rate during three stimulation conditions: stimulation of W1 only (red), W2 only (blue), or W1 and W2 together (green). D, Activity level in the stimulus space, measured as FR and represented by the size of the solid circle. E, Bottom row, activity level in the stimulus space, fitted with the model in (A). The best threshold boundary (red line) maximizes discrimination of the highest frequency (i.e. F1>F2) over the stimulus space. Discrimination is measured as the area under the curve (AUC) using receiver operating characteristic analysis (ROC). Note that multi-whisker suppression (example neuron 2) increases discriminability compared to no suppression (example neuron 1).

### Weighted population averages display reliable frequency categorization across behavioral outcomes

The subjects likely pool activity of multiple sensory neurons to inform its choice so we tracked representation of the two whiskers stimulation intensities at the population level. To do so, the population response Rw1 and Rw2 for the two competing sensory alternatives was modeled as a weighted population average of neurons preferring W1 or W2, respectively (Fig. 5a).

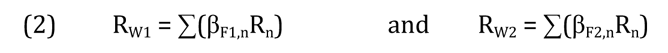

**Figure 5.**
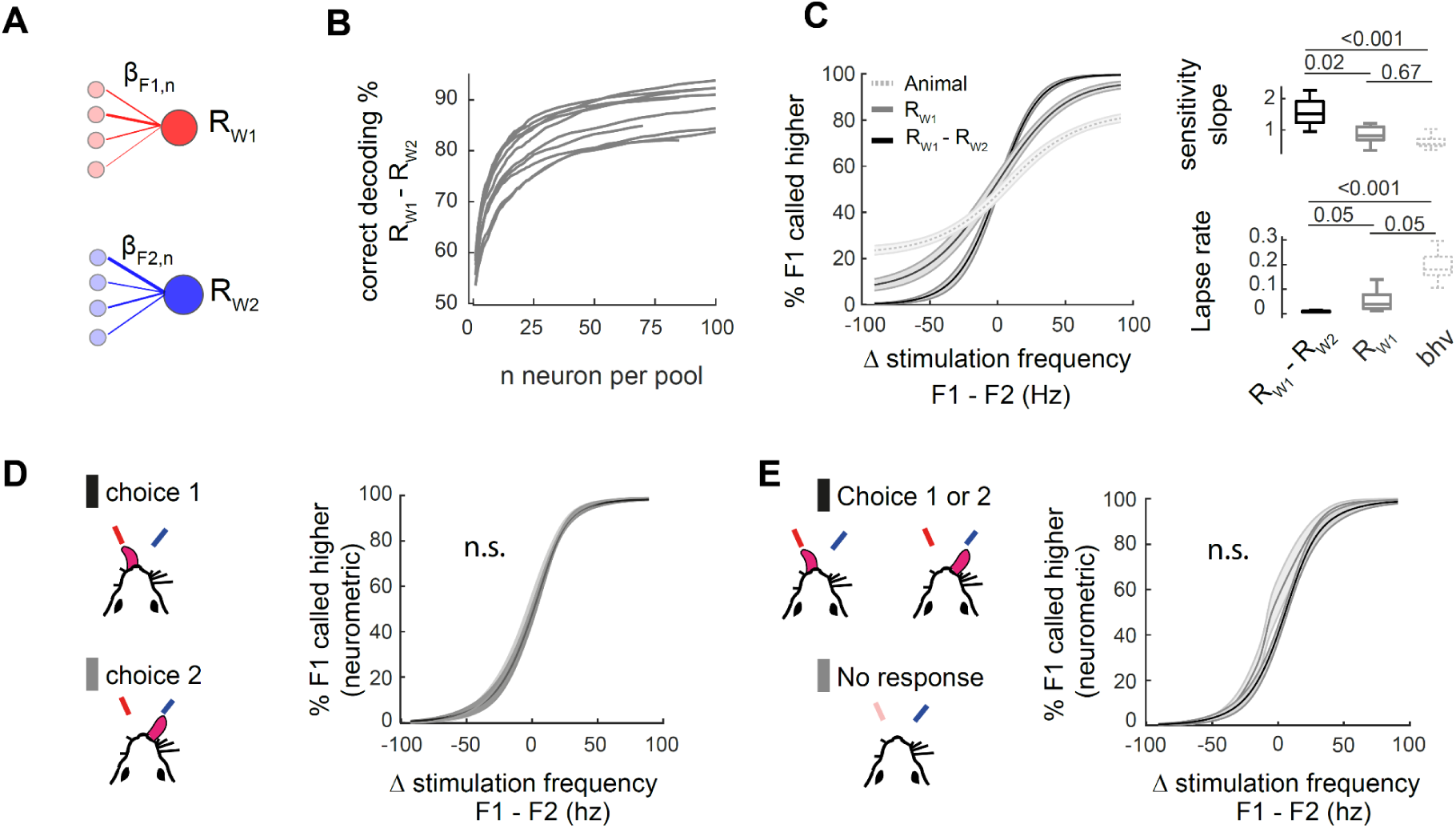
Weighted population averages display reliable frequency categorization across behavioral outcomes. A, Sensory evidence pooled in two sub-populations weighted averages Rw1 and Rw2. Neurons were split in two pools depending on their selectivity for W1 or W2. β weights are obtained directly from the single neuron linear model in equation (1). B, Decoding of the target side (i.e. F1>F2) from the difference Rw1 - Rw2 with an increasing number of neurons in each pool. n = 11 FOVs. C, Neurometric and psychometric functions compared. Left: psychometric/neurometric curves. Right: Comparison of the fitted slope and lapse rate of the three psychometric/neurometric functions. D, Neurometric function of Rw1 - Rw2 computed separately in trials with choice 1 or choice 2. p>0.05 when testing sensitivity slope, lapse rate and bias, n = 11 FOV (Wilcoxon sign-rank test). E, Neurometric function of Rw1 - Rw2 for engaged (black) and disengaged (grey) trials. p>0.05 when testing sensitivity slope, lapse rate and bias, n = 11 FOVs (Wilcoxon sign-rank test).

Where, the coefficients β_F1,n_ and β_F2,n_ from the multiple linear regression were used to weigh activity of single neurons into population averages. By simply thresholding Rw1 and Rw2, we can decode the target side (i.e. is F1 higher or F2 higher), estimating their neurometric performance in the frequency comparison task. This performance can be compared to that of the animals (Fig. 5b-c). When including an increasing number of randomly picked neurons to compute Rw1 and Rw2, we find that integrating information over 6.6 neurons on average (range from 3 to 14; n = 11 FOV) in each sensory pool was sufficient to match the animal’s psychometric performance (same or higher decoded fraction correct; Fig. 5b). When including all sampled neurons, neurometric categorization of Rw1 and Rw2 shows more reliable discrimination than the behavior across trials (lower “lapse rate” for Rw1 and Rw2 neurometric fits; p <0.05, Wilcoxon signed rank test) but have similar sensitivity (slopes compared to the psychometric function; p >0.1). The integrated reading of these two signals by subtraction Rw1 - Rw2 is best solving the task. (p <0.001 slope and lapse rate compared to behavioral performance; Fig. 5c). Overall, these results indicate that neurometric and psychometric functions covary, both as a function of the frequency difference ΔF. Besides, integrating over a small fraction of the population is sufficient to match behavioral performance in the task.

Sensory encoding was then compared across different behavioral outcomes. If the behavioral read-out depends directly on the relative sub-populations firing rate, we should observe a left/right shift in the neurometric functions drawn from trials with left/right choice. To avoid measuring activity related to licks or uninstructed facial movements, we excluded any trials with impulsive movements, i.e. reaction time below the one second stimulation period. We observed no difference in the bias of the neurometric functions of Rw1, Rw2 or Rw1 - Rw2, whether the animal responded left or right (p>0.1, n = 11 FOVs from 7 animals; Wilcoxon sign-rank test; Fig. 5d, Fig. S6). Conversely, the slope and lapse rate of neurometric functions were unchanged whether the animal responded correctly or incorrectly (all p>0.1, n = 11 FOVs from 7 animals, not shown). The same analysis was repeated to compare trials with or without responses (Fig. 5e). Again, the behavioral outcome has no impact on the sensory encoding for frequency categorization (lapse rate, slope, and bias, p>0.1; decoding in engaged versus disengaged trials; Fig. 5e, Fig. S6). A number of different decoding strategies were tested to decode directly the target side, including random tree classifier, bayesian decoder and logistic regression classifier (not shown). In all decoding approaches, frequency categorization performance was unchanged between and error trials (lapse right and slope p>0.05), Conjointly, these results imply that the animal’s perceptual errors are not due to failure in sensory encoding and reciprocally that spontaneous fluctuations in the frequency coding does not affect choice of the animal.

### Sensory and choice coding are orthogonal

We then sought to describe whether and how choice is represented in wS1. Area under the receiver operating curve (AUROC) is a standard metric to test whether a neuron’s activity is correlated to the choice of the subject (termed choice probability or CP(Britten et al. 1993; Crapse and Basso 2015). We selected a matched number of left and right response trials, with small frequency differences (ΔF ≤ 30 Hz), to measure target side discriminability (Fig. 6a) and response side discriminability (CP; Fig. 6b). Across these trials, we find that 22.4% of neurons have different levels of activity depending on the stimulus category F1>F2 or F1<F2 (AUROC different from 0.5; comparison versus bootstrapped distribution with a 95% confidence interval). More specifically, the neurons tuned to W1 have on average an AUROC > 0.5, and neurons tuned to W2 have an AUROC < 0.5, meaning they have higher firing rate distribution when F1>F2, and F1<F2, respectively (mean AUROC = 0.577 and 0.436 for W1 and W2 populations, respectively, p<0.001, Wilcoxon signed rank test; Fig. 6a). If fluctuations in behavioral choice arise from fluctuations in sensory coding, it is expected that populations tuned to W1 and W2 have preferences for the left and right choices respectively. We observe a significant discriminability for the choice side in 20.4% of neurons (CP significantly different from 0.5, comparison versus bootstrapped distribution with a 95% confidence interval; Fig. 6a). Most neurons with significant CP showed preference for left choices (a bias to the contralateral response side, 16.0%, CP >0.5), with the rest preferring right choices. (4.5%, CP<0.5). But how does this choice selectivity relate to sensory selectivity? Considering the population of neurons preferring W1 and W2, we found no difference between the AUROC of these two sub-populations for the response side (mean CP= 0.486 and 0.481 respectively; p>0.1, n= 3706 neurons in total; Mann-Withney U test, Fig. 6b). This analysis indicates that sensory and choice coding exists at the single neuron level and indicates that sensory selectivity does not correlate with choice selectivity.

**Figure 6.**
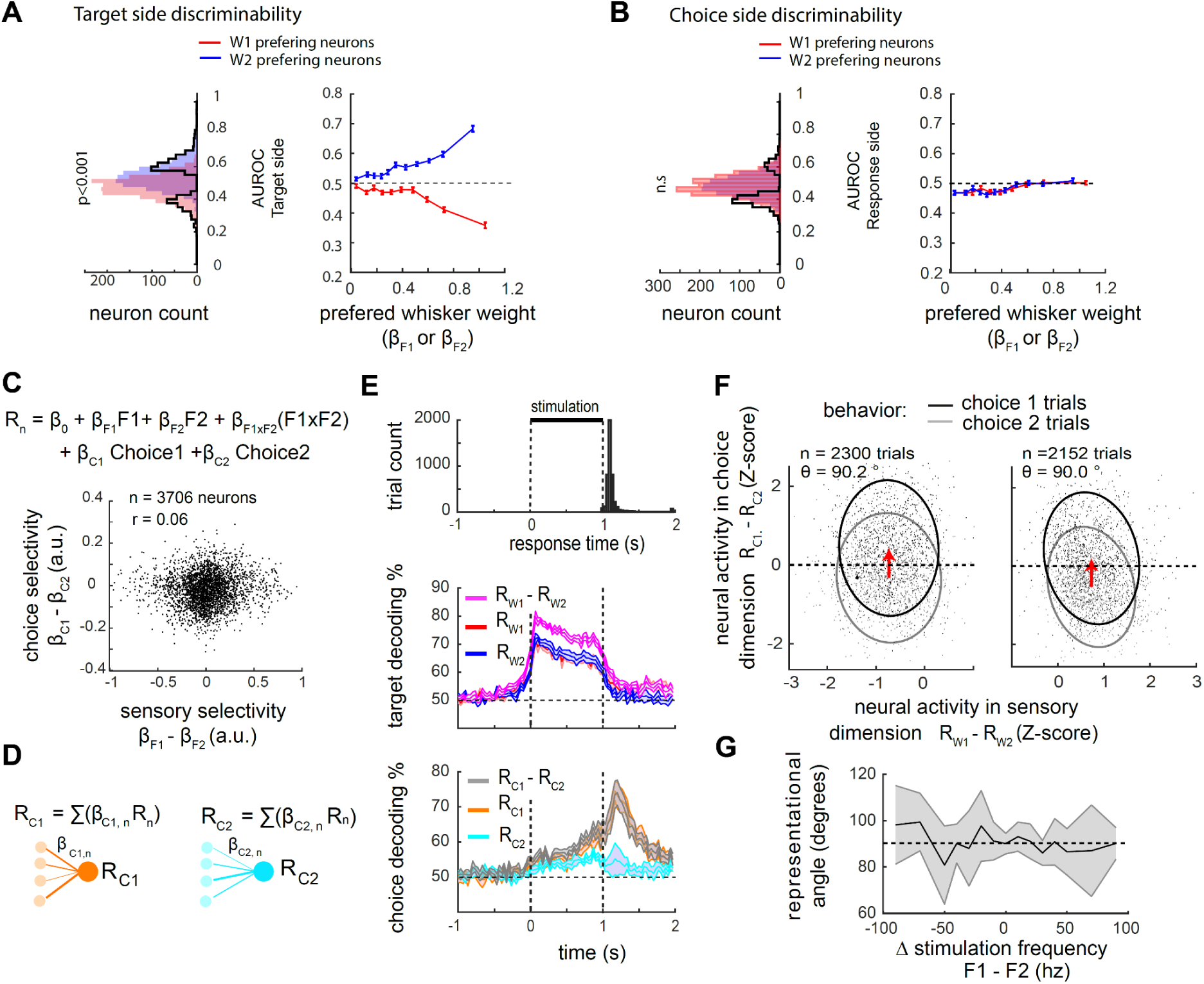
Sensory fluctuations in wS1 are not correlated with behavioral choices. A, Target side discriminability (i.e., F1>F2 versus F2<F1). left; AUROC distribution for all cells (n=3706) black line histogram represent neurons with significant AUROC. right: AUROC as a function of weights for the preferred whisker, averaged in 10 bins containing each the same number of neurons. Only trials with Δf<= 30 Hz and no impulsive movements are included in the analysis. Medians of the two subpopulations are compared with a Mann-Whitney U test. B Response side discriminability (i.e., choice 1 versus 2). Same description as in (A) for choice. Some neurons discriminate choice, as underlined by the black line, but populations preferring W1 or W2 do not show consistent discrimination for choice. C, Top: model including choice 1 and choice 2 as regressor (respectively *β*_C1_ and *β*_C2_). Bottom: weak correlations between choices and sensory regressor relevant to the task (n = 3706 neurons, r = 0.06, p <0.01, Spearman correlation). D, Choice coding evidence pooled into two sub-populations weighted averages Rc1 and Rc2. E, Time course of choice and sensory information in neural activity. Top: Earliest response time of the animals (from video analysis, see Fig. S2). Middle: Prediction of the target side (F1>F2 versus F2>F1) over time, decoding from Rw1, Rw2 or Rw1-Rw2. Bottom: Prediction of the animal’s choice side over time., decoding from Rc1, Rc2 or Rc1-Rc2. Matched number of Choice 1/Choice 2 trials in each stimulus conditions. 10-fold Cross validated. 50% represent chance level. Error shades represent s.e.m. F, Neural activity in the sensory and choice dimensions, in trials with choice 1 (black) or choice 2 (grey). Left: trials with F2>F1 only. Right: trials with F1>F2 only. The red arrow represents transition from choice 1 to choice 2 trials (averages of neural activity across trials). θ is the angle between the x-axis and the transition arrow orientation. Fitted ellipses contain 80% of data points. Choices are best separated on the choice axis, and hardly on the sensory axis. G, Breakdown of the representational angle for the different stimulation conditions. Average across FOVs, only data points with at least 5 FOVs and 5 trials in each FOVs are included. Shaded areas represent s.e.m.

We next questioned to which extent neuronal populations predict the animal’s choice. To do so, left and right choices were included directly as regressors to our multivariate linear model of single cell activity (Fig. 6c; see methods). The distribution of selective regressors weights for choice (*β*_C1_ -*β*_C2_) versus selective weights for whisker frequency (*β*_F1_ -*β*_F2_) display only a weak correlation (pearson’s r = 0.06). This is another indication that neurons selective for whiskers tended to be non-selective for choice and vice versa. The population activity was then averaged by pooling neurons from their weights for left and right choices, yielding two latent variables coding for left and right choices (Rc1 and Rc2, respectively, Fig. 6d), similarly to what was done for computing Rw1 and Rw2 previously. How does the population predict choicesPooling an increasing number of neurons leads to increasing then saturating choice probability in both Rc1 and Rc2 (max CP, 0.62 ± 0.011 and 0.61 ± 0.018, respectively; mean ± s.e.m. n=11 FOV; Fig. S7). Calculation of CP based on the difference between Rc1 and Rc2 is increased when compared to each of these in isolation, and yields a total population max AUROC saturating above around 0.68 (± 0.018; mean ± s.e.m.). We next compared the dynamics of sensory versus choice information (Fig. 6e). Sensory information peaks at stimulus onset and remains high until stimulus offset. Choice information increases slightly above chance at stimulus onset and rise to peak at the response time (Fig. 6e). Thus, decoding of the population reveals reliable coding of choice with ∼68% correct, while decision and sensory information have very different dynamics over the trial time.

The pooling approach allows us to further describe the relationship between sensory and choice encoding with a reduced dimensionality, on a trial by trial basis (Fig. 6f,g). To visualize this relationship, we selected two dimensions of relevance that separate best the stimulus target side and response side respectively (Rw1 - Rw2 and Rc1 - Rc2, Fig. 6f). Overall, we observe that the best separation of left and right choice trials is almost parallel to the sensory axis (Linear discriminant analysis, LDA). A “representational angle” is calculated from the translation of left and right choices in the sensory and choice dimensions. In accordance with our previous observations, choice and sensory representations are encoded orthogonally, (90.2° and 90.0° on average for left and right target trials, n= 2300 and n= 2152 trials; Fig. 6f). This orthogonality generalizes across stimulus conditions and animals (Fig. 6g) and is confirmed for isolated W1 and W2 encoding subpopulations (analysis performed on Rw1 or Rw2 separately, not shown). This orthogonality enables the co-existence of the sensory and choice representations in two distinct dimensions of the neural activity.

### Sensory and engagement coding are mostly orthogonal

How does sensory encoding fluctuate upon different behavioral states? Under the 2AFC design, the absence of behavioral answers (i.e. no licks) cannot lead to a reward. As most no-lick trials occur in blocks at the end of the session, we hypothesize that these represent a distinct state of engagement in the task. We thus sought to characterize some physiological modulation associated with the states of engagement/disengagement. First, a pupillary constriction in engaged versus disengaged trial during the pre-stimulus epoch is observed (p<0.001; spearman’s correlation between no-lick probability and pupil diameter; Fig 7a). Furthermore, engagement was accompanied by a decrease in neural oscillations in the 2 to 10 Hz frequency band (Fig. 7b)(Poulet and Petersen 2008; Jacobs et al. 2020). These pre-stimulus markers strengthen the view of a different brain state during phases of disengagement, in which spontaneous activity of L2/3 is governed by synchronized, large amplitude oscillatory fluctuations of activity.

**Figure 7.**
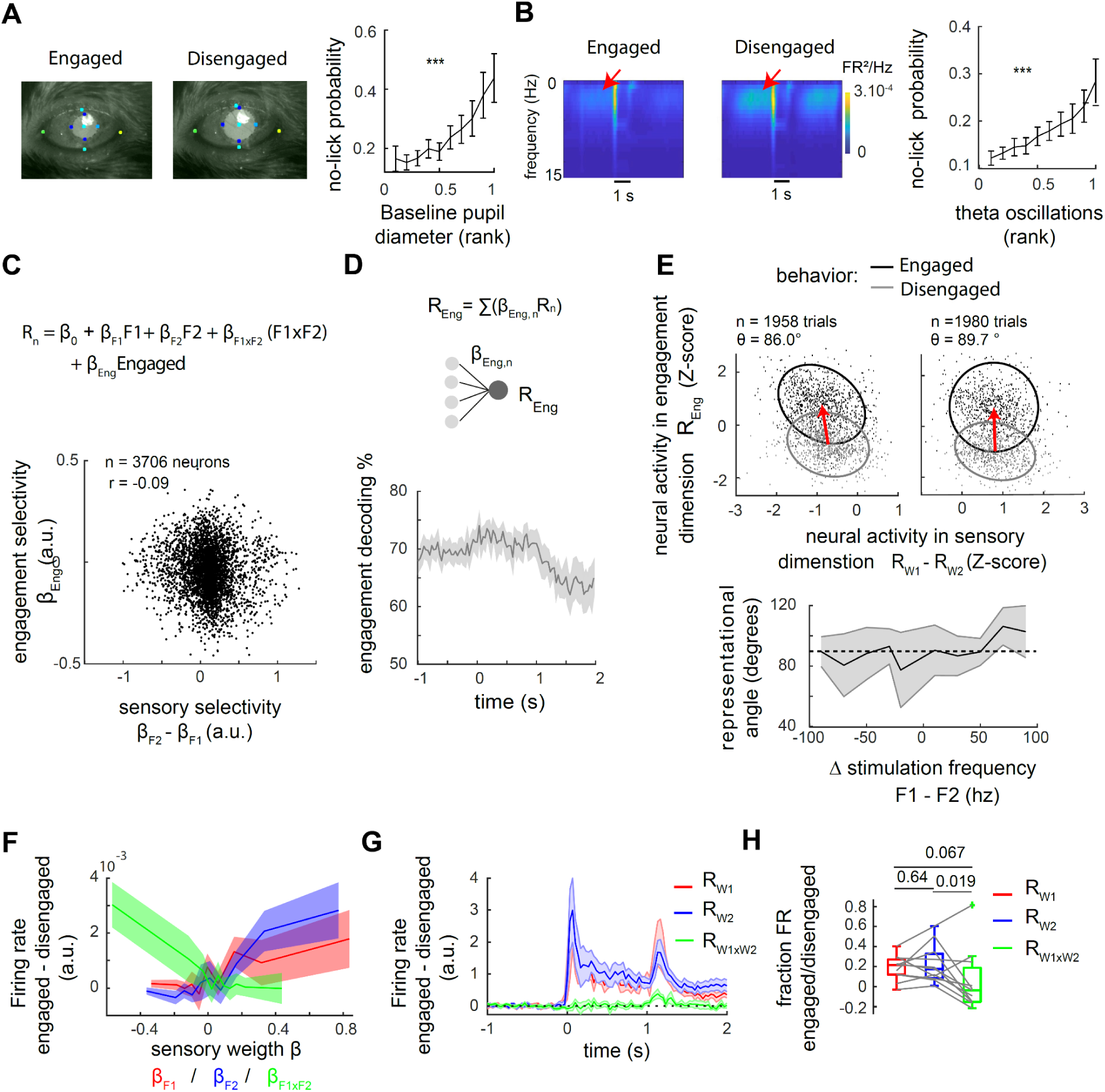
The representation of engagement and whisker evoked activity are mostly orthogonal, with a small gain change. A, Engagement related pupillary contraction prior to stimulus onset. Right: Miss probability in ten deciles with increasing rank of pupil diameter (normalized per session). B, Engagement decreases slow oscillations prior to stimulus onset. Left, spectral power density from one example animal, averaged over all trials. Right, Miss probability in ten deciles with increasing theta power density (normalized per session). C, Top: model including engagement regressor (with weight β_eng_, see methods). Bottom: correlations between engagement and sensory selectivity (n = 3706 neurons, r = -0.09, p <0.01, Spearman correlation). D, Engagement representation over the trial time. Top: engagement coding as a sub-populations weighted averages Reng. Bottom: Decoding of engagement over time. Matched number of trials engaged/disengaged in each stimulus condition, 50% represents chance level. 10-fold cross validated. E, Neural activity during engaged (black) and disengaged (grey) states, in the sensory and engagement dimensions. Left: trials with F2>F1 only. Right: trials with F1>F2 only. The red arrow represents transition from engaged to disengaged (averages of neural activity across trials). θ is the angle between the sensory x-axis and the transition arrow orientation. Fitted ellipses contain 80% of data points. Engagement is best separated on the engagement axis, but hardly on the sensory axis. Bottom: breakdown of the representational angle for the different stimulation conditions. Average across FOVs, only data points with at least 5 FOVs and 5 trials in each FOVs are included. Shaded areas represent s.e.m. F, Engagement related modulation as a function of sensory weights. Computed as the difference in activity in engaged minus disengaged trials (matching stimulus conditions). The neuronal population is split into 10 deciles of beta weights (either β_F1_, β_F2_ or β_F1xF2_) . G, Engagement related modulation of pooled response Rw1, Rw2 and Rw1xw2 over time. Computed as the difference in activity in engaged minus disengaged trials (matching stimulus conditions). Each population is an independent pool of different neurons n = 865, 916, 1925 neurons respectively. Shaded error bars represent s.e.m. across n = 11 FOVs. H, Quantification of the engagement related modulation as the ratio R(engaged)/R(disengaged). statistical comparison across n = 11 FOVs. Wilcoxon rank-signed test.

We then investigated changes in neuronal activity and sensory coding between the two states. To control for movement related activity, we only included trials with a response occurring after the end of the stimulation window. A regressor for engagement was added to the linear model (1) of single neuron activity (Fig. 7c), and the latent variable for engagement (Reng) computed. We found that state modulation in single neurons was little correlated to the whisker selectivity (r = -0.09, n= 3706 neurons; spearman correlation between *β*_Eng_ and *β*_F1_ -*β*_F2_; Fig. 7c). Decoding engagement from weighted average of sub-population preferring engaged versus non-engaged state revealed a significant representation of encoding throughout the duration of the trial (fraction decoded correct = 0.68 during baseline, 0.72 during stimulus presentation; average across FOVs, n= 11 FOVs; Fig. 7d). We next represented neuronal pooled activity in the engagement-related dimension as a function of the neural activity in the task-relevant sensory dimension (Fig. 7d). The shift in activity from disengaged to engaged trials was poorly represented in the sensory discriminative dimension with a representational angle close to 90°, (86.0° and 89.7° on average for left and right target trials, n= 1958 and n= 1980 trials; Fig. 7e). In accordance with our previous observations showing no change in neurometric performance (Fig. 7e and S6), engagement representation lies in a dimension mostly orthogonal to that of the task-relevant sensory representation (Rw1 -Rw2).

However, the analysis for single whisker representation provides a slightly different picture with representational angles differing from orthogonality (93.2 and 98.3 for Rw1 and Rw2 respectively across trials; see details in Fig. S8). In fact, during stimulus presentation, we observe either positive or negative modulation of single neurons firing rate related to the state of engagement, with the average amplitude of single neuron modulation significantly depending on the tuning to whisker frequencies (Fig. 7f). Weights for single whisker frequencies (*β*_F1_ or *β*_F2_) are positively correlated to engagement related gain, while weights for supra-linear whisker interaction (*β*_F1xF2_) of the two whiskers are negatively correlated. Accordingly, the two pooled average Rw1 and Rw2 show increased response amplitude in the engaged state, starting immediately at stimulus onset, while the third pooled average of multi-whisker tuned neurons Rw1xw2 shows a limited decreased response amplitude (fraction of FR engaged/disengaged corresponding to 22% and 14% increase in activity for Rw1 and Rw2 versus a 6% decrease for Rw1xw2; median across n = 11 FOVs. Fig. 7g-h). Engagement thus promote selectively activity related to single whisker versus multi-whisker supralinear activity

## Discussion

Understanding where and how the brain uses sensory inputs to inform perception and behavior is a fundamental question in neuroscience. Here, we developed a 2AFC in which mice had to compare the intensity of stimulation of two whiskers. Few studies required mice to discriminate neighboring whiskers (Ollerenshaw et al. 2014; El-Boustani et al. 2020). This is to our knowledge the first of these discrimination tasks using variation of stimulation intensity, a critical parameter to study perception (Hernández et al. 1997; Stüttgen et al. 2011; Carandini and Churchland 2013). This paradigm takes advantage of the somatotopic map in wS1 to allow manipulations and recordings of two rival sensory alternatives simultaneously.

### A reliable and generalizable neuronal activity substrate in L2/3 neurons for discrimination of neighboring inputs

As described in previous work, we found that evoked activity in L2/3 is somatotopically organized (Woolsey and Van der Loos 1970; Clancy et al. 2015), sparsely distributed in the population (O’Connor et al. 2010; Mayrhofer et al. 2015; Barz et al. 2021) and adapting rapidly in most neurons (Gerdjikov et al. 2018). Increasing stimulation frequency of a single whisker increases monotonically population response (Gerdjikov et al. 2010, 2018; Mayrhofer et al. 2015). Different codes could support perceptual representation of the stimulus, including a rate code and temporal codes (e.g. periodicity of firing)(Recanzone et al. 1992; Johansson and Flanagan 2009). Previous studies have argued that neurometric coding performance by firing rate is closer to psychometric performance than that of temporal coding (Hernández et al. 2000; Luna et al. 2005; Gerdjikov et al. 2010). In our task, the level of activity in its home column is thus a candidate for supporting perception of the whisker stimulation frequency. Our data show a common proportionality between the Δ frequencies, the Δ columnar activation, and the fraction of F1 called higher, suggesting that animals judge frequencies from the relative cortical activation in the two columns (Figure 4).

Besides we observed multi-whisker integration, found to be mostly suppressive (as previously reported in Mirabella et al. 2001; Jacob et al. 2008; Pluta et al. 2017; Laboy-Juárez et al. 2019) with the exception of some neurons responding supralinearly to multi-whisker stimuli (Lyall et al. 2021; Estebanez et al. 2012). Suppression is generally thought to be at least partially cortical dependent (Mirabella et al. 2001; but see Castro-Alamancos 2002) . Suppression is proposed to be beneficial to sharpening boundaries and information coding (Li 1999; Angelucci et al. 2017). In the context of our task, we show that decoding linearly from neurons encoding one single whisker frequency is insufficient to compare frequencies across stimulus space. Rather, multi-whisker suppression provides a more general activity substrate for judging the highest of the two stimuli intensities across the range of stimuli tested (Figure 4).

### Sensory information and representation across states

In our task, the engaged state is associated with a pupil contraction during pre-stimulus period (Ganea et al. 2020), and a suppression of slow oscillatory activity (Jacobs et al. 2020). These characteristics resemble in part to the active mode that displays, when compared to a quiescent awake mode, a depolarization of excitatory L2/3 cells (Crochet and Petersen 2006; Poulet et al. 2012), and a suppression of the large subthreshold oscillations. These effects were shown to be partly under thalamic (Poulet et al. 2012), as well as cholinergic influence (Eggermann et al. 2014; Meir et al. 2018). However, in this active mode, evoked activity is weaker and spatially confined to the principal barrel (Ferezou et al. 2007). Our data show more subtle modulation: we discovered an engagement-related increase of evoked response in a subpopulation coding for single whisker but not in the sub-population encoding for multi-whisker (Figure 7). Thus in the task-engaged state, representation of single versus multi-whisker inputs is promoted, which might favor a more detailed spatial map of the sensory inputs.

However, despite selective sensory gain and some heterogeneity from neuron to neuron, L2/3 populations encoding of frequencies remains stable and whisker selective across states of engagement. Recently, it was shown that brain wide fluctuations of activity was explained by face motion (Musall et al. 2019; Stringer et al. 2019) but also pupil dilation in the awake animal (Stringer et al. 2019). In the later study, motion related activity was found to be encoded orthogonally to that of the sensory coding, but such a relationship was not clear for state only related activity. Here, using cross-validated decoding and excluding face motion from video tracking, we find that task-engagement is reliably represented by L2/3 neurons. Importantly, engagement related activity covary little with variability in sensory encoding from trial-to-trial and is present prior to the stimulus presentation (Figure 7). We thus conclude that correlates of the engagement state are present but mostly in a dimension that lies orthogonally to that of the two whisker’s selectivity. In parallel, we show that the encoding of the frequency comparison information remains reliable across engagement states. This is a first indication that behavioral variability does not stem from alterations of sensory encoding but could originate rather from state related fluctuations in downstream processes.

### Single column activation drives perception of the sensory feature it encodes

Does the animal use wS1 to perform the whisker-intensity discrimination task? Most previous inactivation studies showed impairments of behavioral performance. But impairment of the behavior has several possible explanations: by forming distracting perception for the animal, or through long distance perturbation of neuronal homeostasis (off-target effects)(Otchy et al. 2015; Li et al. 2019). In contrast, our inactivation of focal sources of activity induces specific effects (Figure 3). Alternative methods for manipulation include activation of excitatory neurons or stimulus selective sub ensembles, which were shown to trigger learned behaviors (Salzman et al. 1990; Huber et al. 2008; Sachidhanandam et al. 2013; Musall et al. 2014; Ceballo et al. 2019; Marshel et al. 2019; Buetfering et al. 2022). We believe that both silencing and activating strategies are complementary. Activation studies can delineate a pattern of neural activity sufficient to elicit a behavior. But activation could also trigger perception from an alternative pathway that is not used by the animal under normal sensory circumstances. On the other hand, specific behavioral effects obtained from silencing of an area imply that the area is involved in the sensorimotor pathway that the animal naturally uses. Here we present direct evidence that neural activity in a single column drives conscious perception of the local point or feature it encodes.

The columnar scale has a particular relevance since functional columns are described as building blocks of the mammalian brain, and can be found in almost all brain areas (Mountcastle 1997). Our results apply to early sensory representation in the somatosensory system but could generalize to other sensory representation. In other cortical maps, perhaps more abstractual objects or processes could be manipulated.

From a technical perspective, some limitations of our approaches could be overcome in the future. Activation of inhibitory neurons induces a spread of inhibition larger than the stimulated area (Figure S4). In addition, optogenetic stimulation used here is likely to inhibit activity in deeper layers. L5 neurons may have more impact on perceived orientation tuning in the primary visual cortex, as compared to L2/3 (Marshel et al. 2019). In future studies, holographic activation of single target neurons, in functionally defined ensembles may elucidate further perception related circuits (Packer et al. 2015; Adesnik and Abdeladim 2021).

### Top-down and causal choice signals in the primary somatosensory area

The focus of the study was to investigate the relationship between sensory coding, choice coding, and perceptual report of the animal. In the classical view of perceptually guided decision, sensory evidence builds gradually in decisional and motor areas (Gold and Shadlen 2007; Romo and de Lafuente 2013; Renart and Machens 2014). In this view, noise accumulates over multiple sensory neurons that are weakly correlated with choice. Hence the variability in sensory neurons would explain part of the variability in behavior. Here, we show that the animal’s psychometric sensitivity is matched by a pool of 6 to 7 neurons only and largely outperformed by larger populations (Figure 5), similarly to what was found previously in the somatosensory system (de Lafuente and Romo 2005; Stüttgen and Schwarz 2008). This indicates that mice do not optimally combine the sensory information with the task requirements. It is possible that a source of “noise” is rather added at the level of decision-making circuits downstream of S1. Another possibility -not exclusive with the former-is that read-out of activity by downstream circuits uses a fraction of the population (Buetfering et al. 2022). A feedforward read-out should be associated with a neuronal selectivity for choice, peaking in most ambiguous trials. We found a significant portion of neurons that were indeed selective for choice. However, the encoding of choice across neurons in S1 correlated surprisingly little with the encoding of sensory information. As a consequence, a strong causal contribution of sensory noise onto the variability of the animal’s decision is excluded. The choice signals component begins at stimulus onset, but weakly, and then ramps up slowly until a lick is triggered. Previous studies have indicated that subjects use the earliest temporal component of sensory activity to form their choice (Nienborg and Cumming 2009; Sachidhanandam et al. 2013), while later accumulation of decision signals have little causal influence, and originates most likely from top down modulation (Bondy et al. 2018; Zhao et al. 2020). Our results further show that these later decision signals are mostly unrelated to sensory encoding at the population level. Our results thus favor the model in which perceptual variability does not originate from wS1 but rather from state or choice fluctuations in downstream areas.

The relatively subtle encoding of choice observed here in wS1 is congruent with previous studies of vibrotactile discrimination in non-human primates (Romo and de Lafuente 2013). However, it may seem contradictory with more recent results in the murine model, showing a large difference in evoked activity between hits and misses before the onset of licking in a go/no-go paradigm, and correlation between sensory and choice activity (Kwon et al. 2016; Yang et al. 2016). It is still unclear whether these discrepancies are due to the different animal models, the nature of the perceptual tasks or the different behavioral paradigms. During detection of near threshold stimuli, activity in the primary sensory cortices convey choice signals, in both primates (Palmer et al. 2007; van Vugt et al. 2018), and rodents (Yang et al. 2016). In a situation when animals are asked to perform discrimination of multiple stimuli, the choice might actually be poorly predicted from the primary visual cortex activity (Steinmetz et al. 2019). Therefore, we hypothesize that presence of choice signals in primary sensory areas might depend mostly with the sensory stimulation parameters, with near detection threshold stimuli leading to highest choice predicting activity. In contrast, discrimination of subtle stimuli might be more demanding cognitively, and thus would require more downstream processes for decision making. The comparison of the two vibrotactile objects in our experiments likely falls in the latter case. A distinction shall then be made between detection related signals versus discrimination related signals (Nienborg et al. 2012). In any case future studies would be needed to disentangle better the context of choice signals apparition (Nienborg and Cumming 2014), and their sources (i.e bottom up or top down; (Nienborg and Cumming 2009; Cumming and Nienborg 2016).

The optical techniques presented here allow for a flexible combination of manipulative and correlative approaches, necessary to better understand the substrate of perception and decision making (Panzeri et al. 2017; Stüttgen and Schwarz 2018). The recording of neuronal populations by hundreds enables the investigation of latent choice variables and population coding from trial to trial. We believe that the analysis of the intersection between sensory and choice signals will be essential to understand the features of activity that inform the sensory perception. Here tThese approaches provide insight into the role of primary sensory areas during complex perceptual decisions, namely the formation of a reliable sensory substrate used for downstream decision processes.

## Aknowledgments

We are grateful to Dan Feldman and Gael Moneron for helpful discussions and comments on the manuscripts. We thank Gabriel Lepoussez for providing the VGAT-cre mouse line.

## Authors contribution

P.M.G. and F.H. designed the project and experiments; P.M.G. and S.L.G. performed optogenetic experiments; P.M.G. performed calcium imaging and designed analysis, P.M.G., C.V.R. and A.M. performed analysis; F.H., P.M.G., C.V.R. and D.A.G. designed and built the experimental setup. F.H. acquired fundings and supervised the project; P.M.G. performed visualization and drafted the manuscript with inputs from F.H. All authors provided comments on the manuscript.

## Methods

### Animals

The experiments described in the present work were conducted in accordance with the guidelines on the ethical use of animals from the European Community Council Directive of 24 November 1986 (86/609/EEC) and in accordance with institutional animal welfare guidelines and were approved by Animalerie Centrale, Médecine du Travail and the Ethics Committee CETEA of Institut Pasteur, protocol numbers.

All mice were aged 8 to 16 weeks at the time of surgery. 7 C57BL/6J male mice were obtained from Charles River. 2 GAD-67 and 9 VGAT-cre male mice were bred in house Animalerie Centrale, Institut Pasteur. All animals were kept under a 12h light-dark cycle, with food available *ad libitum*. At the end of the experiment mice were aged at maximum of 32 weeks

### Surgery

Mice were anesthetized with isoflurane (Piramal critical care, UK; induction: 4%, maintenance: 2%) and placed in a stereotactic frame (Kopf Instruments, USA). Mice were injected with buprenorphine (Vetergesic CEVA, France; 0.02 mg/kg, subcutaneous.) for pain management and their eyes were protected from desiccation by applying ointment. The fur was removed over the skull, the skin was disinfected with betadine (Mylan, USA), and xylocaine (Bayer, Germany; 0.25%, 0.05 ml) was injected subcutaneously for local analgesia. The skull was exposed and several whisker barrels (at least 3 barrels among one of these: rows A-E within arcs 1-2, and/or alpha beta gamma) in the right hemisphere were identified using intrinsic optical imaging together with whisker stimulation by means of piezo control (Mayrhofer et al. 2015). A 3 mm diameter round craniotomy was then performed and a virus was injected at 350 µm depth from the pia, with multiple injection spots (from 200 to 400 nl injected in each) patterned in a grid injection spaced by 500 µm, such that viral expression would span the entire window. The virus carried either a red calcium indicator alone (7 mice; jRGECO1a; AAV1.Syn.NES.jRGECO1a.WPRE.SV40; a gift from Douglas Kim & GENIE Project (Addgene plasmid # 100854 ; http://n2t.net/addgene:100854 ; RRID:Addgene_100854) (Dana et al. 2016), or in combination with EGFP (2 mice pAAV.synP.DIO.EGFP.WPRE.hGH, 1/20 dilution, viruses mixed prior to the injection). For the optogenetic, the virus carried either Channelrhodopsin 2 alone (5 mice, pAAV-EF1a-doubleloxed-hChR2(H134R)-EYFP-WPRE-HGHpA; a gift from Karl Deisseroth (Addgene viral prep # 20298-AAV9 ; http://n2t.net/addgene:2029) or with a green calcium indicator (4 mice; pAAV-EF1a-double floxed-hChR2(H134R)-mCherry-WPRE-HGHpA; a gift from Karl Deisseroth (Addgene viral prep # 20297-AAV9 ; http://n2t.net/addgene:2029 with pAAV.CAG.Flex.GCaMP6s.WPRE.SV40; a gift from Douglas Kim & GENIE Project (Addgene viral prep # 100842-AAV9; http://n2t.net/addgene:100842; 1/10 dilution (Chen et al. 2013). Injections were performed using beveled glass pipettes (Drummond, USA) with a diameter of 15-30 µm. The craniotomy was then sealed with glass coverslip (3 mm diameter round window; UQG Optics Ltd, UK). Dental cement (DE Healthcare Products, UK) was applied to keep the window in place and to form a head cap holding a custom-made head-post made of titanium. Throughout surgery, body temperature was maintained at 37 °C with a feedback-controlled heating pad (Thorlabs, USA). After surgery, mice received carprofen in a gel (Dietgel, clear H2O, USA) for pain management (0.02 mg/kg; every 24 h; 3 d postoperatively). Mice were single-housed for the rest of the experimental procedures to avoid potential damage to the implant.

### Behavioral task

Water deprivation of the animals started a minimum of 2 weeks following surgery to allow for recovery. Animals were handled twice a day with 1 ml of water delivered manually by the experimenter through a syringe and progressively habituated to head fixation and to the experimental setup. This procedure has been described in detail in a previous article (Mayrhofer et al. 2013). Once the animals were habituated to the setup, all whiskers but two were trimmed below 3 mm, on the opposite side to the craniotomy. The pair of target whiskers were neighboring on the same arc: either B1/C1, or C1/D1 were picked. At the beginning of each session, the two target whiskers were inserted in two capillary glasses mounted on two independent piezo elements set at 3 mm from the whisker pad. Animals were first exposed to stimulation and got free water delivered 500 ms after onset of the stimulus. Water was delivered according to the contingency decided as follow: higher whisker on the arc (either B1 or C1) was associated with left water spout and lower whisker on the arc (either C1 or D1) with the right water. The percentage of water delivered automatically was progressively lowered (∼10% per day), and thus water was only rewarded when the animal performed a lick on the correct side. To avoid frustration during this learning phase, droplets of water were sometimes added manually on the correct side, via a direct command on the custom software, after the animal produced licks to the incorrect side. Progressively, animals learned the contingency, and performed better over the course of days/weeks with a variable learning rate (Fig. 1). Once this association was mastered by the animal (>70% correct responses in at least two consecutive days), simultaneous stimulation of the two whiskers was progressively introduced at a low rate. Once the performance was maintained at a high level on a simultaneous whisker stimulation task (p<0.05 assessed by a chi² compared to chance level, test across discrimination conditions), we started the imaging on a daily basis; Image collections were performed throughout a period of less than one month.

### Psychometric/Neurometric analysis

Evaluation of the performance of the animal/classification of neuronal signals was done by quantification of parameters of a psychometric/neurometric function. A logistic function of the following form was fitted: (Whichman and Hills 2001):

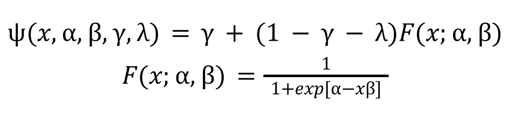

*γ* and *λ* correspond respectively to the lower and higher asymptote of the fit. α the steepness of the curve, and β the value of the midpoint, or bias (where fractions of left and right licks are equal). All parameters of the fit were left free to vary. Comparison of psychometrics function were performed as the comparison of best fit parameters between groups of mice or FOV (e.g., comparison of α, the best fit in engaged versus disengaged states with a Wilcoxon signed rank test).

### Behavioral video data analysis

Images of the animals’ faces were acquired by video tracking at two possible rates: 60 Hz (mice 1-3) or 300Hz (mice 3-7). Images were processed with the DeepLabCut toolbox (Mathis et al. 2018) at the frame rate of acquisition. For the video tracking from below the snout, we first tagged 400 frames picked randomly across all datasets with the following tag: nose left, nose right, nose center, philtrum, teeth center, chin, tongue, front right whisker base, front right whisker tip, back right whisker base, back right whisker tip, left paw, and right paw. The neural network was trained to detect the markers on a dataset of 500 frames randomly selected among all videos. Through visual inspection, another 200 frames with unsatisfying tagging were manually labeled and added to the training dataset; the network was then re-trained with all labeled frames. All videos were processed with the latter trained network. The marker positions measured in pixels and the likelihood of detection provided by DLC software were used to compute the movement of the different body parts. In each session, we computed standard deviation of x-y position of the nose and whisker markers (in pixels). These measures were ranked and averaged for each trial. We then identified the 5% trials with most and 5% trials with least face movements, 5_high_ (so trials with high movements), and 5_low_ (trials with the least movement) respectively. The idea behind this strategy stems from the observation that in every session there are trials with high levels of face motion and trials in which the animal remains still. These trials are in our view the most reliable way to normalize datasets across sessions and imaging parameters. From this strategy it is possible to compute relative values of face, snout and whisker movement within the session, as used throughout our study, although these are not given as absolute measures in degree angle or millimeters.

To estimate tongue movements, we used the baseline epoch from the 5_low_ trials to define a threshold, computed as average plus three standard deviations of the likelihood of appearance. For instance, tongue was considered as outside of the mouth when its likelihood at a given time point was superior to the above described threshold. When detected, tongue movements were classified as left or right depending on the marker relative x position compared to teeth (fixed, computed as the average of all frames within the session). The whisker angle was first computed as the average of all 4 whisker markers in y dimension. Nose position was first computed as the average between the three nose markers in the X dimension. Both nose and whisker measures were Z-scored session per session, as follows:

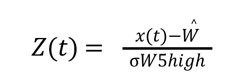

With *W* being the average position during baseline across trials, and σW_5high_ being the standard deviation of position across the 5_high_ trials. From several possible normalization procedures tested, this normalization provide most comparable distribution histograms of relative nose and whisker position across sessions. Finally, to compute the earliest reaction time, we used the Z-scored nose position. If the nose position changed by more than 0.1 Z-score between two frames, the first of the two frames was counted as the reaction time.

The same labeling procedure was used on videos acquired from the side of the face (all recorded at 30Hz from 4 animals), with the following set of markers: eyelid top, eyelid bottom, pupil left pupil top, pupil right, pupil bottom, pupil center. The pupil diameter was computed as the Euclidean distance between markers ‘pupil left’ and ‘pupil right’ markers. When likelihood of detection of any of these two markers was below 0.5, the trials were thrown away. From a measure in pixels, pupil diameter was normalized between the 5th and 95th percentile for each session separately as follows: P(t) = (P(t) - P5th)/ (P95th - P5th). This measure yields a relative change in pupil size as compared to the dynamic range it can achieve, with 90% of the values between 0 and 1. The use of low/high percentile was preferred to min/max because of possible outliers.

### Two-photon calcium imaging data acquisition

Two-photon imaging was carried out after a minimum of 2 weeks following surgery to allow for sufficient viral expression. Functional images were acquired at ∼29.7 Hz using bidirectional scanning and an image resolution of 512×512 pixels spanning 738 x 605 microns. The field of view was centered using an optical imaging signal (Fig. 2a) in order to cover two barrels of whiskers within the same FOV. The genetically encoded calcium indicator was excited with a Ti:sapphire laser (jRGECO1a at 1040 nm, GCaMP6s at 920 nm; pulses frequency of 80 MHz, Chameleon; Coherent) using an equivalent power of ∼100 mW for jRGECO1a and a power ranging from 25-120 mW depending on the depth of recording for GCaMP6s. Emitted light was recorded through a 16x, 0.8 NA objective (Nikon, Japan)and detected with two photomultiplier modules having the following filter settings: band pass filter 540/40 and 617/73 for green and red respectively and short pass filter BrightLine 750/sp for each channel (Semrock, USA). two-photon laser scanning was controlled using the ScanImage software (Pologruto et al. 2003).

### Simultaneous two-photon imaging and optogenetic stimulation

Simultaneous optogenetic stimulation and calcium data acquisition was performed in the same setup as standard two-photon calcium imaging with the following differences. Images were acquired continuously in 5 planes, at a total volume imaging rate of 4.58 Hz. Optogenetic stimuli consisted in a train of 1ms pulses delivered at 40 Hz from -0.25 s before the onset of the stimulus to 1 s after the offset of the stimulus. Blue light pulse trains were generated with a LED (470 nm, M470L4, Thorlabs, driver LEDD1B, Thorlabs) taking continuous voltage from a breakout box (National Instruments, SCB-68A) as driving input. From the light source, we mounted serially a diffuser, a lens, and an iris was positioned such that its image is formed in the first imaging plane. The iris’ image diameter was set to either 0.10 or 0.45 mm for the selective and broad inhibition respectively. This setup only allowed us to perform the three optogenetic conditions described in Fig. 3 in the same two photon areas, but in different sessions. To do so the imaging planes were first matched and the image of the iris was then positioned using a XY translation mount. Trials with different light intensity (0 to 44.3 mW.mm² for selective and 0 to 17.9 mW.mm² for broad light patterns) were randomly alternated with trials without light. We used a Polychroic mirror (Chroma, zt470/561/nirtrans) transmitting the infrared and blue light, while reflecting the green to the gated PMTs (Hamammatsu, H11706). The triggerable PMTs shutter was synchronized with each pulse of blue light and engaged during 1.33 ms. For each pulse of blue light, a fraction of the frame was missing and thus interpolated from previous and following frames, on a line-by-line basis (Matlab interp2 function, linear interpolation). Because we observed a decay of neuropil fluorescence stemming from the blue light pulse, the interpolation was carried over a total duration of 10 ms per light pulse (i.e. roughly one third of the imaging frame). Two-photon imaging data then followed the same processing pipeline as standard two-photon imaging.

### Optogenetic inhibition during behavior

A projector (DLP4500, Texas instruments, USA) was synchronized with the behavioral software. Light intensities were matched to calibrations made in the simultaneous two-photon and optogenetic. 40 Hz trains of 1 ms pulses were used. The surface of the projected disk was set to 0.105 mm for the selective inhibition (using from 8.3 to 35.9 mW.mm², measured under constant light) and 0.45 mm or 0.64 mm for the broad inhibition (5.0 to 21.8 mW.mm²). In the 3 animals for which individual calibration was available we used an individualized light power ranging from 8.3 to 19.4 mW.mm². In other animals which only expressed channelrhodopsin, we chose a light power of 19.4 mW.mm². We did not observe changes as a function of light power applied (not shown), thus behavior in trials across light power were pooled together. For each individual animal, a set of images corresponding to the three stimulation conditions was generated upstream from behavior, based on the intrinsic optical imaging data. Every behavioral session started with the positioning of the system using a manual translation mount. The image of the projector was focused ∼150 µm below the brain surface. Trials with and without optogenetic were randomly intermingled with an average proportion of roughly one out of four trials with light stimulation on a brain location, and three out of four on the dental cement, i.e. sham condition.

### Two-photon calcium imaging data preprocessing

In 11 FOV from 7 animals with jRGECO1a, we carried detailed analysis over complete population statistics. Movies were motion corrected and ROIs delineated using the routine from suite2p (Pachitariu et al. 2016) ROIs were manually curated and every visible cell with an event rate > 0.005 Hz was included. The mean ROI fluorescence was corrected by local neuropil, and then by subtracting a 30 s rolling 10th percentile (to exclude slow drifts). Neuropil alpha subtraction factor was computed independently for each ROI as the slope of a linear regression of neuronal fluorescence against neuropil fluorescence (Runyan et al. 2017). The regression was carried out only outside of activity epochs as defined by the fluorescence being below the 16th percentile of the entire time-series (putative period of non-activity). More than 95% neuropil correction factors computed this way were between 0.5 and 0.85, values below or above this range were respectively set to these boundaries. Finally single ROI fluorescence were Z-scored and a constrained deconvolution was applied (Pachitariu et al. 2016) to return a continuous spiking estimate. This preprocessing strategy and criteria were optimized from the freely available jRGECO1a dataset from the CRCNS website (Dana et al. 2016, Mohar et al. 2016) to match the state of the art algorithmic performance in event detection (Berens et al. 2018).

### Linear regressions and sensory evidence / choice side reconstruction

We used a linear model to regress the recorded spiking activity of single neurons against the series of different stimulation applied to the two whiskers. Models were fitted separately for each neuron. Spiking activity was averaged In all trials over a 1 second epochs during both intertrial intervals (1 sec before stimulus onset) and during whisker stimulation (0 to 1 sec after stimulus onset). All epochs of activity were concatenated into a single vector Y that was regressed against the square root of the two whisker frequencies (F_1_ and F_2_) and the product of these two regressors (F_1×2_).

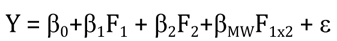

In the versions of the model that explicitly included choice (Figure 6) or engagement (Figure 7), we used only delayed response or no response trials, so that activity related to face movement did not interfere with the analysis. We used two binary regressor separately for left and right choice (set to NaN during baseline epochs) set to 0 or 1 depending on the choice in the ongoing trials:

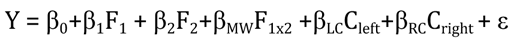

To model engagement, we included a single regressor that was set to 0 in non-engaged trials or 1 in engaged trials, in both baseline and during stimulation. Non-engaged trials were defined as the animal not licking a spout and occurring in the last third of the daily session. Engaged trials were defined trials with a response from the animal and occurring in the first half of the session:

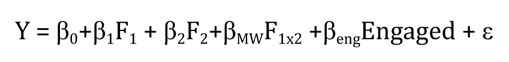

Before regression was actually applied, we normalized the activity of single neurons by dividing it by its average value. This normalization enables the comparison of weights across cells as they would be proportional to the variance explained by the regressor. With the aim of comparing the weights between regressors, we also z-scored the regressors values. establishing thus a direct proportionality between regressors weights and activity level. The model fitting was performed using matlab fitlm function that uses a QR decomposition algorithm. Later, reconstruction of sensory evidence is computed separately for each regressor as the sum of weighted neuronal activity divided by the sum of weights. Pooled activity summaries (Rw1, Rw2,…) are computed from independant pools of neurons, by choosing only cells tuned to the summarized signals more than to the other alternative (e.g. pooling only neurons preferring w1 to compute Rw1, for instance). The result of this computation is a relative estimate of whisker identity strength. In figure 6 and 7 however, all neurons were included to compute the task-related variables. Another decoding strategy would be to reconstruct the frequency via maximum likelihood estimation using bayesian inference (Runyan et al. 2017). However, we chose linear pooling for its simplicity as it provides an intuitive way of summarizing activity over pools of neurons and does not rely on any assumption of independence between neurona’s activity.

