## Supplementary figures for "Coexistence of state, choice, and sensory integration coding in barrel cortex LII/III"

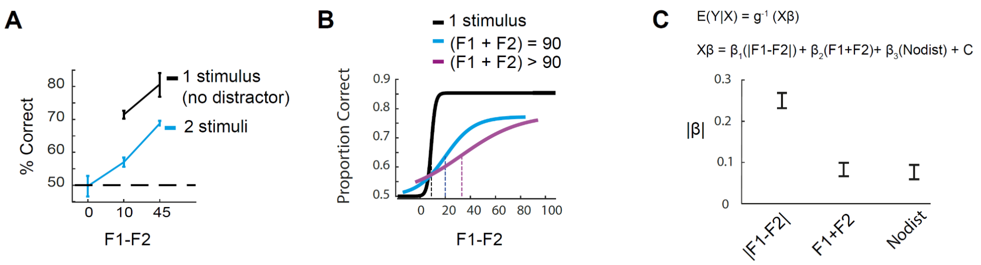


**Figure S1. Behavioral performance in the stimulus space**

A, Average performance across trials with single (black) or dual (blue) frequency stimulation. In details: at $\Delta F = 0$, F1/F2 is equal to 45/45 (blue). At $\Delta F = 10$, F1/F2 is equal to 50/40(blue) or 10/0 (black). At $\Delta F = 45$, F1/F2 is equal to 45/0 (black) or 90/45 (blue).
B, Psychometric fit of the proportion correct P(correct) for different categories of stimuli. Vertical bar represents sensitivity threshold at P(correct) = 0.65. Discrimination is easiest when a single stimulus is presented. When two stimuli are presented on W1 and W2 simultaneously, discrimination is harder when the sum of the two frequency is high (F1 + F2 >90Hz)
C, The F1-F2 difference is the most important stimulus parameter for behavioral categorization. Influence of the stimulation parameters on the subjects' performance (average across pooled trials from all experiments, n=166599 trials). The response boolean vector Y (success =1, failure =0) is fitted with a generalized linear model, assuming a binomial distribution and using a logit link function. All three regression variables were Z-scored so that their **β** weight are comparable. Note that we compare absolute **β** weight. C is a constant. Error bars represent a 95% confidence interval.

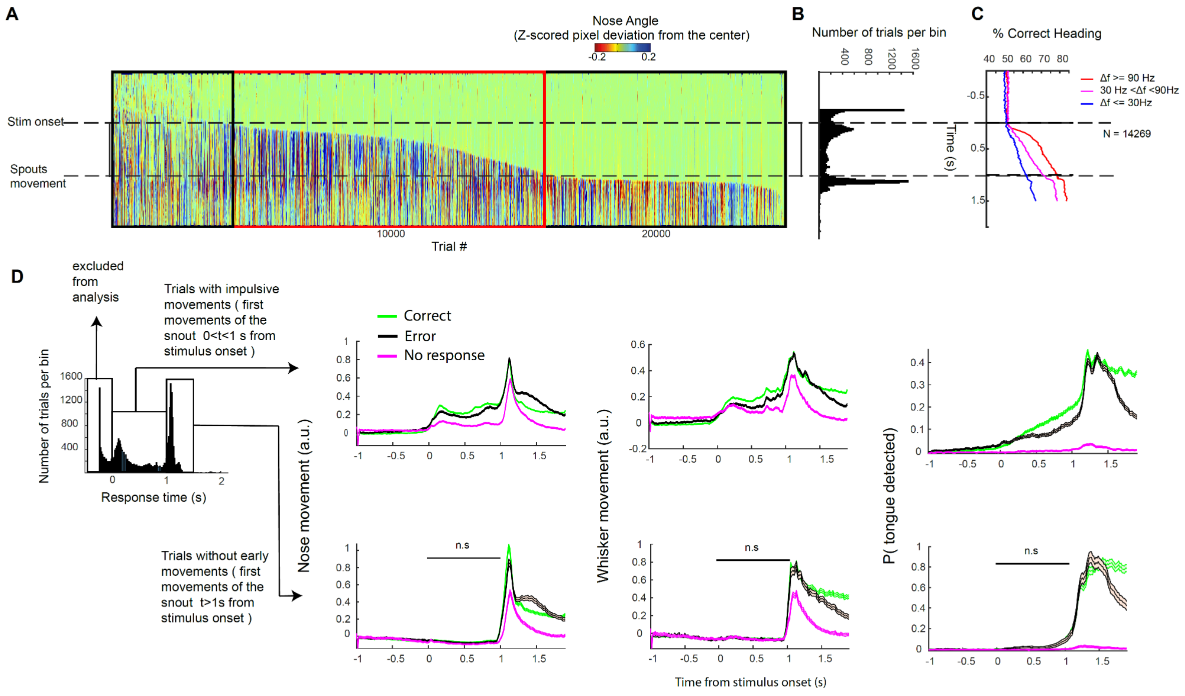


**Figure S2, Reaction times and facial movements**

A, Nose heading direction as a function of time. Trial # in x, time in y, blue indicates left heading nose; red indicates right heading nose; trials sorted according to the first detected nose movement (see methods).
B, Distribution over time of first detected nose movement.
C, Heading direction predicts the correct target side for easy (red), middle (magenta) and difficult trials (blue).
D, Body movement extracted from video analysis (see methods). Left: trial sorting according to nose reaction time. Right, top row: body movement during impulsive trials. Right, bottom row: body movements during late response trials. p-value indicates difference of movement between the three behavioral categories (Friedman test, n = 7 animals). Error shades represent a 95% confidence interval following a normal distribution on a trial-by-trial basis (n= 30300 trials from 7 animals).

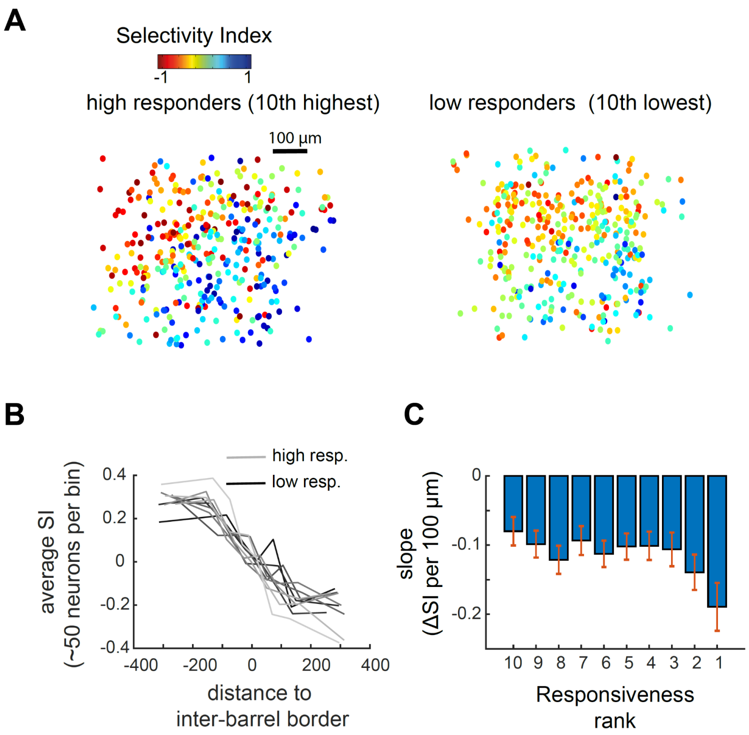


**Figure S3. Highly responsive neurons are more whisker selective and clustered toward the barrel center.**

A, Spatial distribution of the selectivity from highest and lowest responding neurons (left and right respectively).

B, SI as a function of distance to the two-barrel separation depends on responsiveness of the neurons. From lightest to darkest color line represents neurons split in ten deciles going from the most to least responsive neurons. Data is further split in bins of ~50 microns width and finally averaged to allow plotting the spatial dependence of SI.

C, Slopes from a linear regression of the data in B, between distance to barrel and selectivity. Selectivity of highly responsive cells (Responsiveness rank 1) changes more rapidly with distance to the barrel center. Error bars represent CI95.

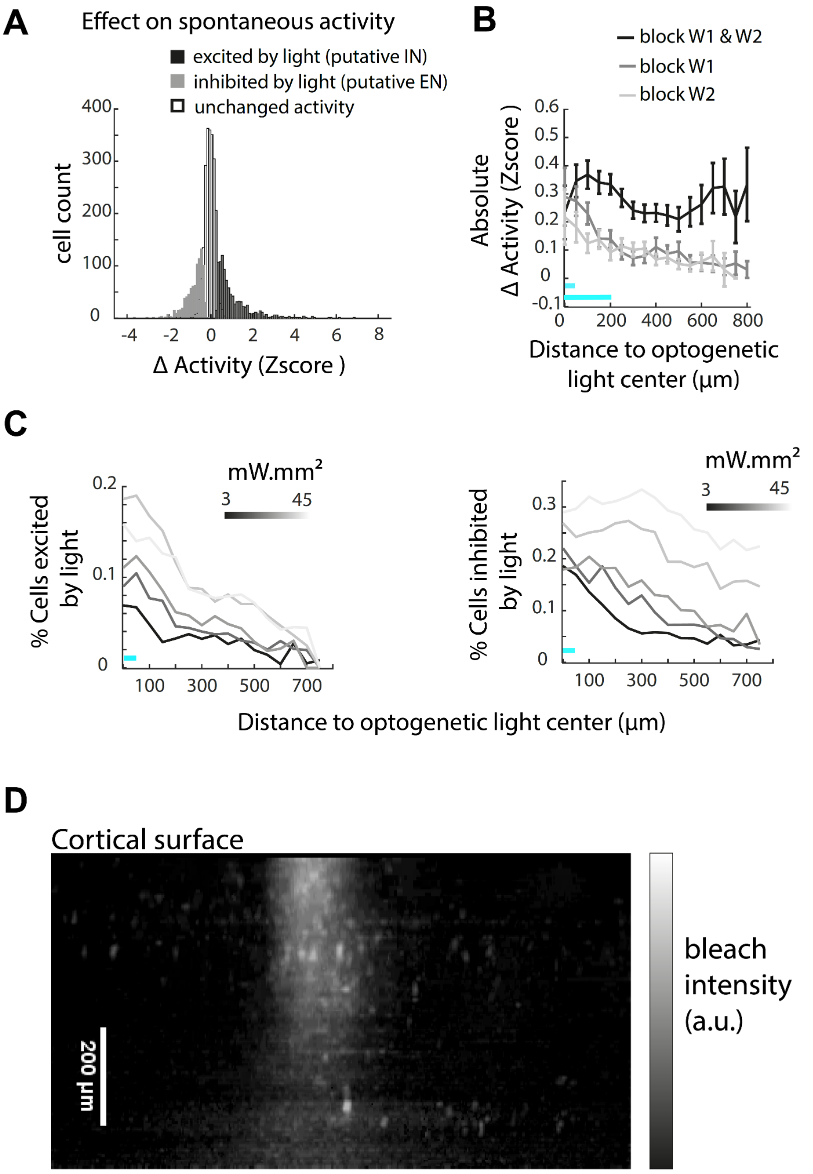


**Figure S4. Spatial spread of optogenetic excitation/ inhibition**

A, Change in fluorescence in response to optogenetic light (450 µm disk) for all cells in the FOV ( n= 5034 neurons, 3 animals, 5 FOV ). 23.2% of cells showed activation by optogenetic light (putative inhibitory neurons) and 23.8% cells showed inhibition (putative excitatory neurons) compared to spontaneous activity.

B, Change in activity as a function of distance to center of inhibition, quantified as absolute change in activity |ΔZscore|, for the three optogenetic conditions with the smallest light amplitude (e = 1.4 $mW.mm^2$ and e= 3 $mW.mm^2$ for the 450 µm and 105 µm disks respectively).

C, Inhibition and excitation as a function of distance to illumination center during selective barrel illumination.

D, Visualization of the optogenetic light spread in the axial dimension of the microscope. The light spread is measured as bleaching induced on GCaMP6s expressed across cortical layers. Bleaching was induced by prolonged exposure to the optogenetic light pattern (12 minutes of continuous illumination, 105 µm disk FWHM, at ~ 44 $mW/mm^2$). Pixel intensity is computed as fluorescence before exposition minus fluorescence after exposition, thus bleaching appearing as a brighter column. Fluorescence was measured before and after bleaching with the same two photon imaging parameters in a stack with 5µm steps. Each plane is averaged over 1 second. Volume rotation and visualization was performed with ImageJ.

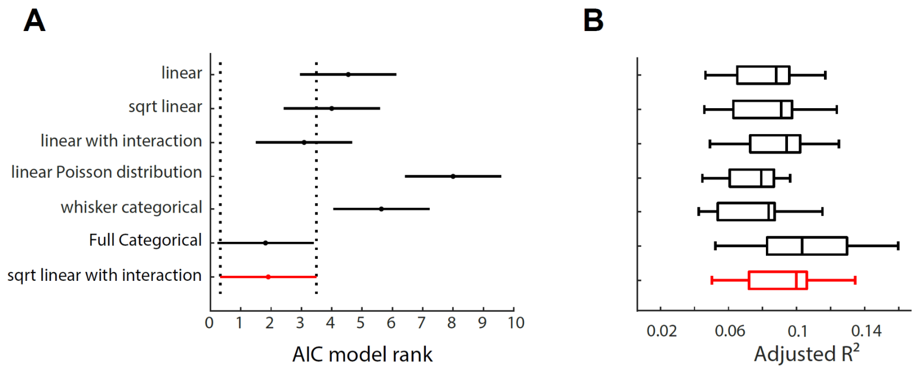


**Figure S5. The square root linear model with interaction yields similar results to the full categorical model for predicting single neuron's activity.**

A, Comparison of Akaike criterion (fit performance given complexity of the model). Friedman's test with multiple comparison, shows that no other model tested outperform the model used in the rest of the study (sqrt linear with interaction; described in Equation (1); see methods)

B, Variance explained for different linear, categorical, and non-linear models of activity. The higher variance explained by the full categorical model could indicate that some neurons have a modal tuning to specific frequencies.

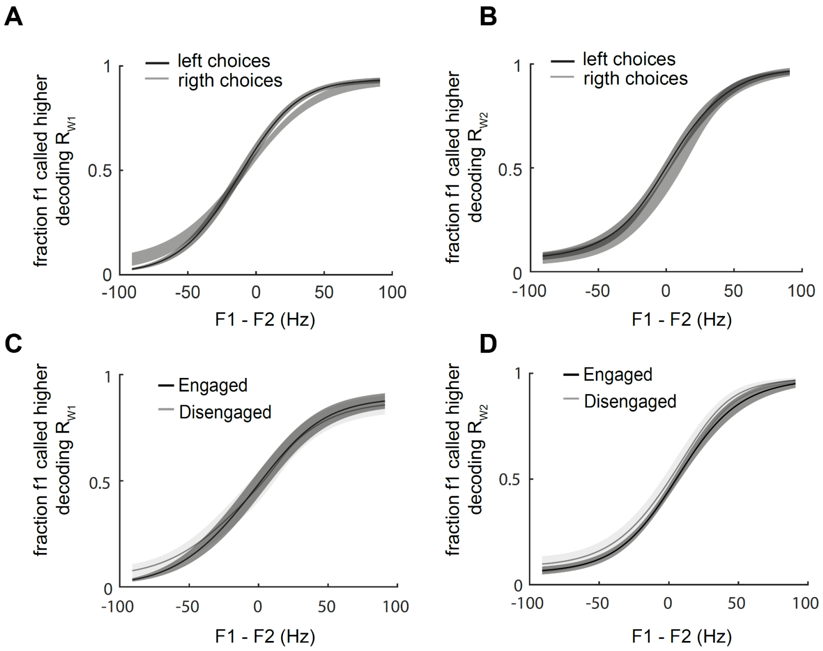


**Figure S6**: **Neurometric functions of Rw1 and Rw2**

A, B, Neurometric functions in trials with left versus right response from the animal. Decoding of Rw1 (A) or RW2 (B).

C, D, Neurometric functions in trials with or without responses. Comparison of slope and bias compared with a Wilcoxon sign-rank test reveals no significant difference ($p>0.05$, n = 11 FOVs). Decoding of Rw1 (C) or RW2 (D).

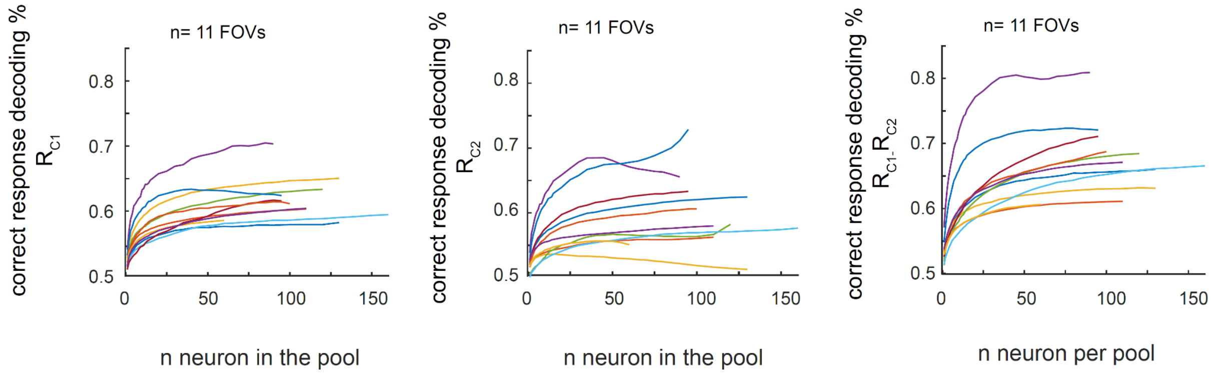


**Figure S7 choice information in sub-populations $R_{C1}$, $R_{C2}$ and $R_{C1}$ - $R_{C2}$.**
Performance of choice discriminability as a function of the number of neurons included in the pooled response. From left to right: discriminability of $R_{C1}$ ,$R_{C2,}$ and $R_{C1}$ - $R_{C2}$. Each line represents data from one FOV (n = 11 FOV). Matched number of Choice 1/Choice 2 trials in each stimulus conditions. 10 fold Cross validated.

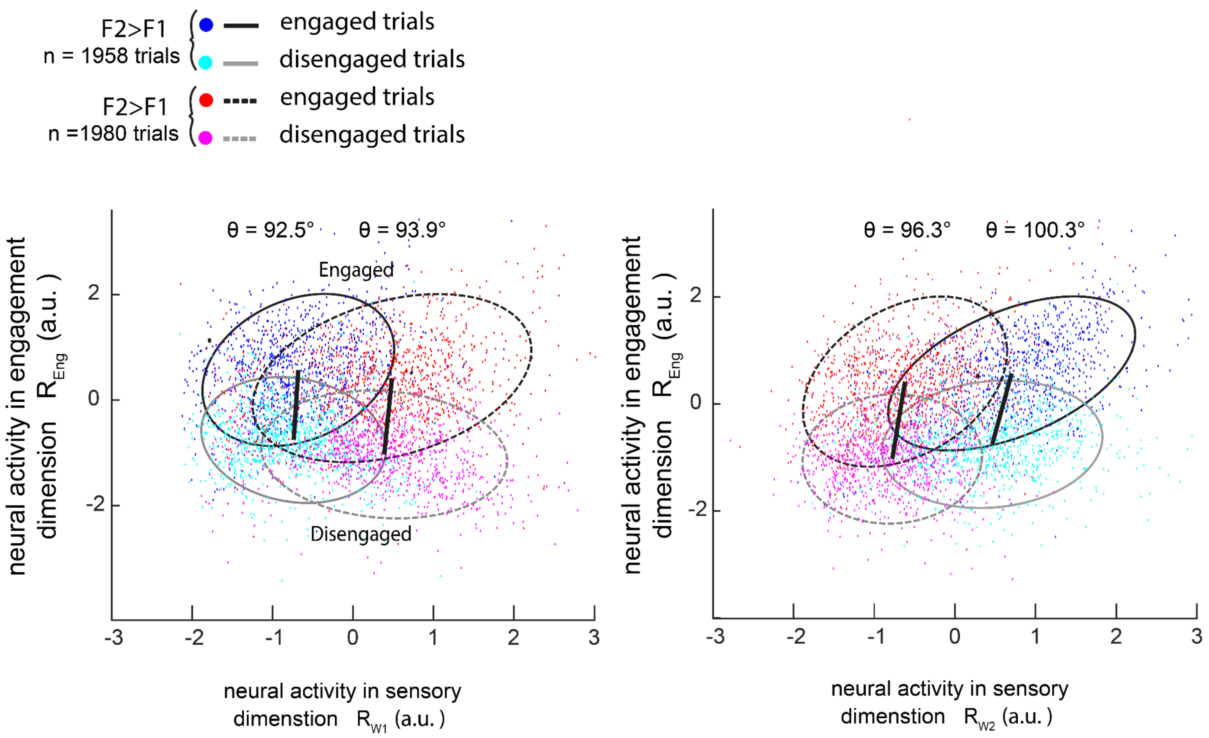


**Figure S8. Engagement changes the sensory gain for the preferred and non-preferred whisker.**

A, Neural activity during engaged and disengaged states, in the sensory and engagement dimensions. Left: engagement versus W1 neural representation (Rw1). Right: engagement versus W2 neural representation (Rw2). Same method and description as in Fig. 7e. but both F1>F2 and F2>F1 trials are included in the same panel. Bars represent transition from engaged and disengaged trials. Note that all representational angles are tilted to the right, showing an increase in response of Rw1 and Rw2 during engagement across stimulation conditions (i.e. independent of the whisker stimulated at the highest frequency).
